# Single-nucleus RNA sequencing revealed the impact of post-mortem interval on the cellular component and gene expression analysis of mouse brains

**DOI:** 10.1101/2024.12.30.630832

**Authors:** Yunxia Guo, Junjie Ma, Xiaoying Ma, Kaiqiang Ye, Chao Jiang, Jitao Xu, Yan Huang, Xi Yang, Qinyu Ge, Jianyou Zhang, Guangzhong Wang, Hao Huang, Xiangwei Zhao

## Abstract

Accurate analysis of cell atlas and gene expression in biological tissues using single-nucleus RNA sequencing (snRNA-seq) is dependent on the quality of source material, and post-mortem interval (PMI) is one of the major sources of variation in RNA quality. Although the use of RNA-degraded tissues in transcriptome analysis remains controversial, such samples are sometimes the sole means to address specific questions. Current studies on the impact of PMI on transcriptome data are limited to large-scale RNA-seq, which ignores cellular heterogeneity. Thus, deciphering the non-cell-autonomous effects caused by PMI is imperative for understanding the cellular and molecular disruption it elicits. Here, we investigated the impact of PMI on cellular components and gene expression using snRNA-seq data from mouse brain tissues of post-mortem. We collected samples that were allowed to decay for varying amounts of time at 25°C prior to snRNA-seq, covering the entire range of RIN values. The different effects on the PMI to the degradation rate of mRNA and rRNA within nuclei, and the mRNA presented a more stable state. Multi-channel analysis revealed the preferential transient depletion oligodendrocytes and OPCs with increasing PMI. In addition, a rapid widespread overregulation of ribosomal transient recruitment to protein (RP) genes in various cells, and reached a plateau at PMI of 36h. Although state depletion of neuronal cells was not detected, we reported significant upregulation of PMI-dependent RP genes in its subpopulations and their cell loss. Moreover, RP genes showed the greatest differential expression in the subpopulations with greater cell perturbation, and we speculated that aberrant expression of these genes might be associated with cell death. In this study, we systematically investigated the changes in the transcriptome profile of brain tissue induced by PMI at single-cell resolution, and revealed one of the important factors that might be responsible for the changes. In addition, our data complemented a possible explanation for the changes in the cellular state of brain tissue induced by postmortem hypoxia-ischemia, and provided a reference for transcriptome studies of RNA degradation samples.

## 1. Introduction

Post-mortem biological tissues are a valuable resource for biological research and a key resource for studying gene expression patterns, especially for brain tissues. Specifically, the use of post-mortem material is crucial for studying the patterns of normal gene expression underlying tissue specificity within individuals, as sampling such tissues from living individuals would be impossible. Death does not imply that the billions of cells in a body have ceased functioning, but rather that they are no longer working together. After cessation of blood flow or similar ischaemic exposures, deleterious molecular cascades commence in mammalian cells, eventually leading to their death [1]. Connections between cells are not break down, but what happens to these cells in the hours or even days at the end of life is unclear. In addition, the death of an organism triggers a cascade of events that ultimately, in a relatively short time frame, lead to cell death and autolysis. Although DNA is known to be relatively stable over long post-mortem periods, RNA is much more labile in nature. One of the important factors for RNA fragmentation is postmortem time, and the processes triggered by the post-mortem interval (PMI) can significantly alter physiologically normal RNA levels. Currently, there are conflicting reports on how the post-mortem interval affects RNA integrity. The effect of PMI varies widely across different tissues [2–6], and brain tissues are most significantly affected by postmortem hypoxia-ischemia [7]. In addition, the responsiveness of different cell types to PMI should also be inconsistent. However, these results have been detected by traditional bulk samples transcriptome or immunohistochemical techniques [8–10].

Single cell// nucleus RNA sequencing (sc/snRNA-seq) has revolutionized the ability of researchers to explore cellular heterogeneity and genetic variation at the single cell level, and has become a suitable technical means to analyze cellular networks. However, the technique remains controversial with respect to the use of RNA-degraded biological samples. It has been shown that sequencing samples with low RNA quality (as measured by the RNA integrity index [RIN]) results in lower quality of the obtained transcriptome data [11–13]. In addition, the current commercial sc/snRNA-seq technology platform, based on poly-A capture, requires a RIN requirement of samples greater than seven, but the inability to ensure high-quality preservation of samples under specific conditions leads to RNA degradation, which forces degraded samples to become the preferred tissue to solve specific problems. Moreover, the samples collection in the field or in clinical settings, in which extracted RNA is likely to degrade and may not faithfully represent in vivo gene expression levels[14]. Therefore, it is important to explore whether the severely degraded RNA samples can be subjected to high-throughput snRNA sequencing and to investigate the effects of postmortem time on cell composition and gene expression.

Mouse postmortem brain tissue is an important resource for studying gene expression changes in neurological diseases. However, transcriptomic studies of such tissues rely on the integrity of mRNA molecules during PMI periods. Here, we mimicked the effect of PMI on transcriptional data at room temperature by storing postmortem mouse brain tissues at 25°C in an enzyme-free environment for different periods (PMI = 0 h, 24 h, 36 h, 48 h, 54 h), and snRNA-seq was performed on these samples to gain insight into PMI-induced cell composition and gene expression changes of brain tissues. This study provides the first comprehensive and detailed exploration of the molecular changes of degraded brain tissue caused by PMI, which opens the horizon for further transcriptomics study of degraded samples, and may also provide reference for transcriptome study of degraded FFPE samples.

## 2. Results

### 2.1 Effect of PMI on mRNA quality in mouse hippocampus

To evaluate the effect of PMI on mRNA quality, we implemented a mouse hippocampus model. The isolated hippocampus was subjected to ischemic injury under a constant temperature (25℃) without enzyme, including five groups: a flash-frozen control group (H0 with minimal/0 h PMI) corresponding to four experimental groups with distinct PMI (24h: H24, 36h: H36, 48h: H48, 54h: H54). The RIN (RNA integrity number) values of these tissues decreased significantly (*p* < 0.05) with prolonged PMI, spanning almost the entire RIN range (from 9 to 3), indicating severe RNA degradation (Fig. S1A, S1B). To explore the effect of PMI on RNA molecules at the single-nucleus scale, we extracted nuclei from all groups of samples and performed high-throughput single nuclear nucleic acid amplification (10× Genomics). Unexpectedly, the main peak size of cDNA of nuclei across all samples was about 1000 bp (Fig. S1C and Fig. S2A), fully meeting the requirements of commercial 3’ library preparation, indicating that PMI negatively affects mRNA in hippocampal nuclei less than total RNA. Although the cDNA fragments distribution was excellent, since they were sorted and purified by AMPure XP magnetic beads, the resulting yield will directly reflect the mRNA responsiveness to PMI. As we expected, the cDNA yield of the H24-54 groups decreased by nearly 20 to 40% compared to H0 (Fig. S1D), indicating that the corresponding proportion of mRNA was fragmented and discarded at the time of cDNA purification. But the main peak of the library after the same amount of cDNA input was 450-500 bp, and the yield was similar (Fig. S2B). Overall, PMI had a smaller negative effect on mRNA than total RNA in the nuclei, or was more sensitive to RNA within the cytoplasm. Our findings demonstrate that snRNA-seq can be successfully performed on mouse hippocampus samples with a PMI of up to 54 hours, breaking the legend that severely degraded samples cannot be used for transcriptome sequencing.

### 2.2 Non-significantly effect of PMI on snRNA-seq quality control data

RIN value and the cDNA quality could only reflect the degree of mRNA fragmentation and loss within the nuclei, not its detailed transcription level. Therefore, we next analyzed the effect of PMI on snRNA-seq quality control data. We calculated exon inclusion levels, and the results showed that the proportion of exons gradually increased with the prolongation of PMI, although not significantly (Fig. S3A-D), which is consistent with our previous results of paraformaldehyde-fixed samples [15]. In eukaryotic cells, introns will gradually accumulate to maintain normal cell growth during the growth attenuation stage caused by nutrient deprivation [16, 17], contrary to our conclusion. It has also been reported that fresh human brains possess more introns than postmortem samples, possibly indicating that a large number of pre-mRNA transcripts may be involved in RNA editing [18]. Therefore, the regulatory role of introns in organisms under external persecution needs to be further explored. We then calculated the number of effectively detected nuclei using the same analysis strategy, under the premise that the number of nuclei entered into the snRNA-seq platform for each sample was comparable. A greater effect of PMI on the nuclei numbers was observed, with approximately 50% less nuclei in H54 than H0 (Fig. S3F). Therefore, with the same sequencing throughput, the sequencing saturation of H24-H54 samples naturally increased (Fig. S3E), and the median of genes and UMIs per nuclei were also increased gradually, although not significantly (*p* > 0.05) (Fig. 3G, H), and the intra-group repeatability was high (Fig. S3J, K). The analysis of gene detection sensitivity showed no association between sensitivity and PMI (Fig. S3L). In addition, the percent of mitochondrial genes also showed an increasing trend with the extension of PMI (Fig. S3M). We speculate that energy changes in brain tissue during postmortem time cause an increase in the proportion of mitochondrial genes. Housekeeping genes are a class of genes that can be stably expressed in all cells, and their products are necessary for maintaining the basic life activities of cells. Such as tubulin genes, glycolytic enzyme genes and ribosomal protein genes. The expression level of housekeeping genes is less affected by environmental factors and is continuously expressed in most growth stages or almost all tissues of an individual, often in the euchromatin of the nucleus. We were curious whether such stable housekeeping genes were affected by PMI, so we further explored the effect of PMI on the expression of some housekeeping genes (*Tpm1*, *Alb*, *Actb*, *Gapdh*, *Hprt*, *Ppia*, *Bhmt*, *Srp72*, *Mylk*) and found that the expression of most housekeeping genes was up-regulated, although also not significant (*p* < 0.05) (Fig. S3I). This is consistent with the results of postmortem human brain [18] and similar to the pattern of housekeeping gene expression changes in brain tissue after fixation with PFA [19]. In general, PMI had no significant effect (*p* > 0.05) on the quality control results of snRNA-seq sequencing, which could be used for subsequent analysis.

### 2.3 snRNA-seq substantiates PMI-dependent deficiency of oligodendrocytes and OPCs

The effect of prolonged hypoxic-ischemia on the cell types of mouse brain, although reported, have been unclear. To explore the effect and mechanisms of PMI induced cell composition of mouse brain at the single-cell resolution, and monitored their changes over a period of five points. we performed and integrated snRNA-seq to build cellular maps of 8-week-old mouse hippocampus with different PMI (Fig. 1A). After filtering out potential doublets, and poorly sequenced and damaged nuclei, a total of 123,464 individual nuclei were arranged by uniform manifold approximation and projection (UMAP) in 2 dimensions for visualization (Fig. 1B). Unsupervised clustering revealed largely similar cellular landscapes across all sample groups (Fig. S4A, B), and a total of 14 distinct clusters in these samples were identified (Fig. 1B, C). These clusters were manually identified eight major cell types based on the basis of the expression of known cell-type-specific markers [20–22] as excitatory neurons (Ex, cluster 0, 1, 2, 6, 7, 10, 12), inhibitory neurons (Inh, cluster 8), astrocytes (Ast, cluster 3), oligodendrocytes (Oligo, cluster 4), microglia (Micro, Cluster 5), Endothelial (Endo, Cluster 9), oligodendrocyte progenitor cells (OPCs, Cluster 11) and cajal-retzius cell (CRC, cluster 13) (Fig. 1B, C and Fig. S4C, D). Then we counted the proportion of the main eight types of cells, and found that the nuclei of Ex accounted for the largest proportion (69.9%), followed by AST (8.2%), Oligo (6.8%), Micro (5.3%), etc. (Fig. 1D). Except that the proportion of Oligo cells was less than half of that reported in the literature, the frequency result of each cell types across all samples were generally consistent with the cell proportions of mice brain [21]. In addition, containing few nuclei, Cluster 13 was omitted from further analyses.

**Figure 1.**
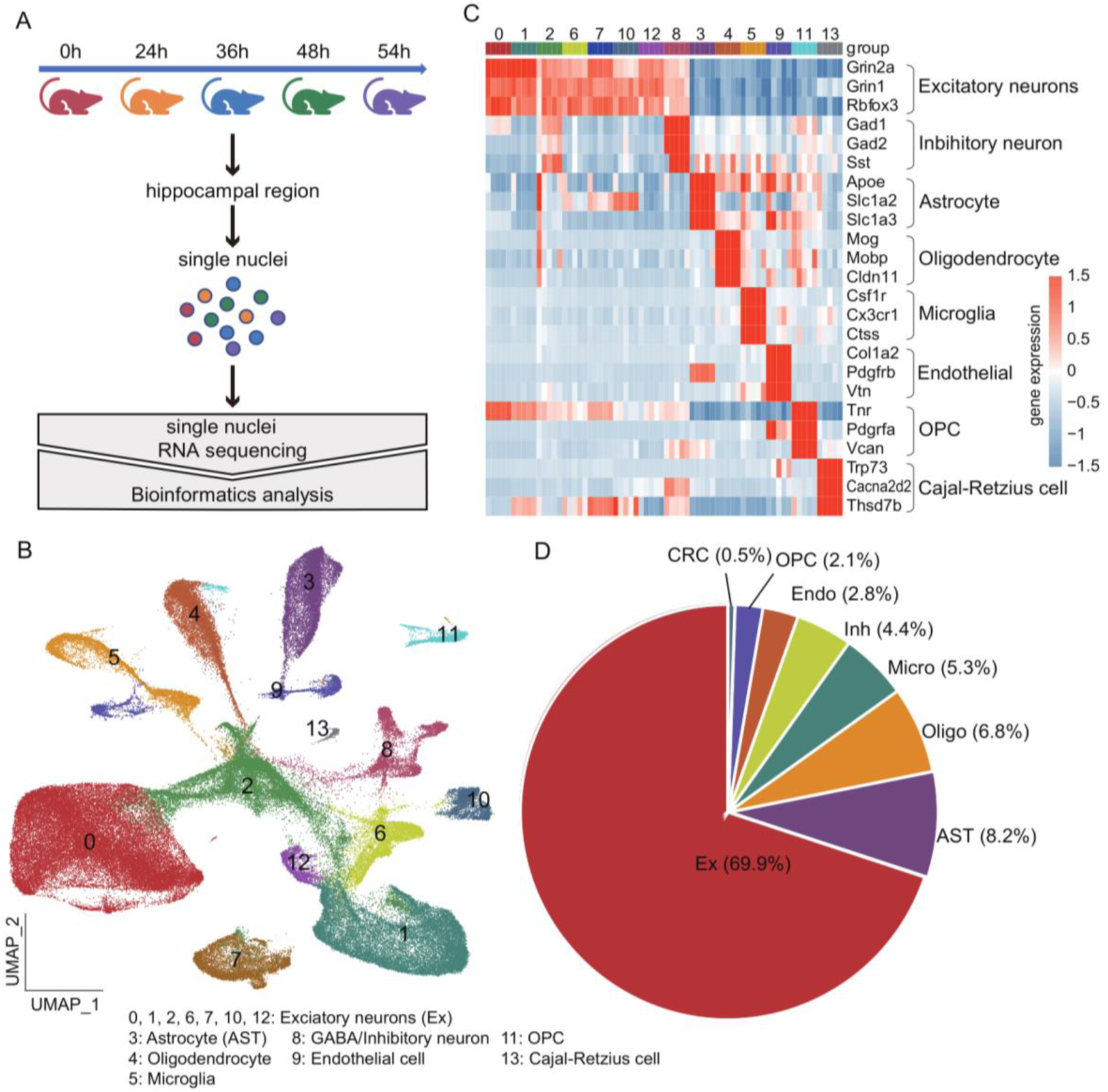
snRNA-seq distinguishes major cell types in hippocampal region from different degraded samples. (**A**) Overview of study design. (**B**) Uniform manifold approximation and projection (UMAP) layout showing major cell types. UMAP plot showing 14 distinguished clusters, 123,464 total nuclei, with cell-type identities as determined by expression of specific markers. (**C**) Heatmap representing expression of specific markers in every sample, identifying each cluster in B. (**D**) Pie chart shows the frequency of each cluster across all samples. n = 3 biologically independent mouse brain samples per group.

To compare the responsiveness of cell types to PMI based on transcriptomic changes across all experimental groups, multi-dimensional approaches were used. Most intuitively, UMAP plots directly reflected the similar distribution of nuclei from the five post-mortem samples (Fig. 2A), indicating that all cell types can be detected. However, the nuclei were sparsely distributed in the H24-54 groups (Fig. 2A), which was related to the number of cells detected. We then examined changes in the composition of each cluster in the context of PMI. Compared with H0, most clusters were similarly represented in H24-H54 groups, but an increase of Ex at H24 that was exaggerated at H36 (Fig. 2B), and the exception of Ex was overrepresented by about 15 %. The Oligo and OPCs were underrepresented with a cliff pattern, and these changes began at H24 and reached the bottom at H36 (Fig. 2B). In contrast, other glial cells representation was stable throughout the PMI. However, Oligo and OPCs diminution coincided with increase of Ex cells, the direct cell ratio only showed a reciprocal change, and we could not determine whether the increase of Ex percent was caused by the loss of Oligo and OPCs. To more accurately identify the cell types that responded specifically to the PMI, relative frequency analysis of clusters in each group was performed to avoid cell proportion fluctuations within the sample (Fig. 2C). Again, the cell state changes of Oligo and OPCs were the most obvious (Fig. 2C), further evidence that the diminution of Oligo and OPCs was partially PMI dependent. Moreover, the correlation between changes in other cells and PMI was minimal and insignificant (*p* > 0.05), but stood out of those in Oligo and OPCs cells (Fig. 2D). Finally, to compare cell type responsiveness to PMI based on transcriptomic changes across experimental groups, we performed Augur prioritization [23] and highlighted cell-types with the greatest transcriptomic divergence (Fig. S5A). Prominent change in Oligo was also identified (*p* < 0.05), consistent with cells validated in above results and prompting detailed subanalysis. Micro and Endo cells were less affected by PMI, consistent with the results reported for ischemic pig brains [24]. However, inconsistent with the results obtained in the warm ischemic porcine brain, PMI had little effect on Ex cells in our study, which may be termed “delayed neuronal death” [8] or perhaps related to differences in postmortem brain processing. Comparisons between all experimental groups and the H0 revealed significant enrichment of gene sets enrichment of facilitating cytoskeletal assembly, axonogenesis, regulation of signal transduction by p53 class mediator, axon ensheathment, glutamatergic synapse-related, and other cell necroptosis pathways were significantly revealed significant across Oligo cells in the brains investigated (Fig. S5B, Table S1).The results indicate that Oligo cells undergo a series of biological processes such as inflammatory death, necrotizing death and glutamate poisoning during postmortem.

**Figure 2.**
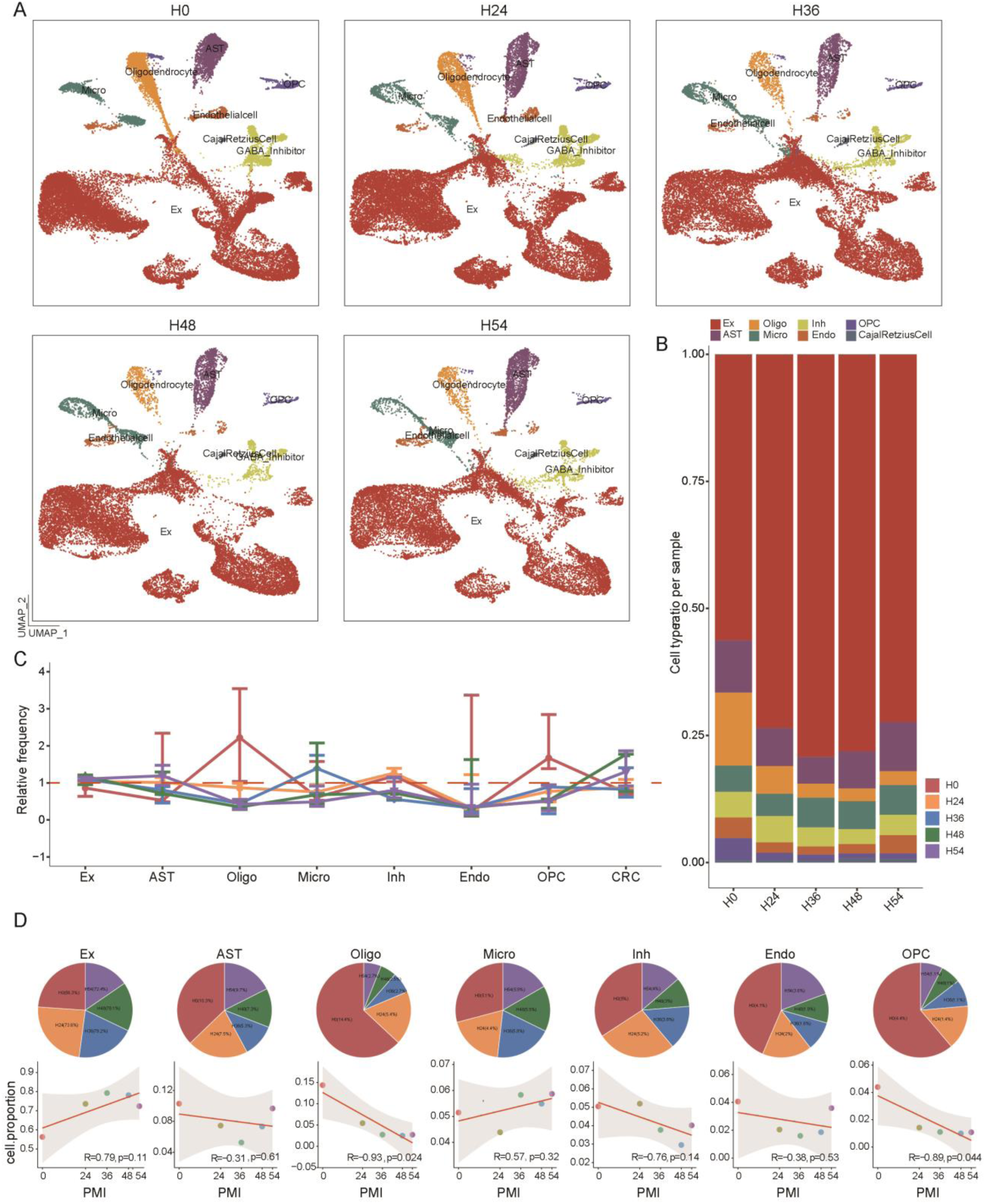
Cell-type-specific transcriptomic changes assessed by snRNA-seq across various PMI. (A) UMAP plots of distribution of nuclei of different cell types at PMI of 0h, 24, 36h, 48h and 54h. See also Figure 1B and Figure S3C for the distribution of cell-specific markers. (B) Bar charts depicting the frequency of representation of cell types in every sample. (C) Relative frequency of clusters in each sample, normalized to overall frequency in Figure.1D. (D) Pie chart and scatter plots showing frequency and the correlation with time for each cluster across each sample. Cluster 13 had very low frequency and were omitted from downstream analysis. n = 3 biologically independent mouse brain samples per group.

Taken together, these observations indicate that, all cell types could be detected in brain tissue after 24h to 54h of postmortem ischemia, and most cells had similar proportions, which was unexpected. Nevertheless, we ultimately identify the cell types most significantly affected by PMI: Oligo cells and OPCs, which will be the focus of further research.

### 2.4 Similarity of gene expression pattern driven by PMI

To reveal the effect of PMI on gene expression in different cell types, we next focused on differentially expressed genes (DEGs) on a per-cell types basis, specifically comparing datasets from H24-54 versus H0 groups. A significantly (adjust *p* < 0.05) overexpressed co-gene set was found, with the genes encoding ribosomes, misfolded proteins and mitochondria, such as *Cmss1*, *Lars2*, and *Ubb* or stress genes, independent of cell types and sample groups (Fig. S6). The fold change (log2FC) of these significantly up-regulated genes varied in different cell types and showed an increasing trend with the prolongation of PMI (Fig. S6). We speculate that there may be a gene set whose expression changes are independent of cell type and only related to PMI. Next, we screened genes with the most significant changes in the 28 comparison groups to identify the functions of these PMI-related genes. We expected to set a threshold to identify co-occurring DEGs (co-DEGs). We counted the times of co-DEGs that detected in 28 comparison groups, and drew the frequency distribution density plots as a reference for setting a reasonable threshold (Fig. S7A, B, top). We selected DEGs as co-DEGs as long as they appeared in the 10 comparison groups, and 213 co-up DEGs and 267 co-down DEGs were screened respectively (Fig. S7A, B, bottom). The co-upDEGs were mainly enriched in translation, ribosome, protein binding (ubiquitin-protein ligase, protein kinase, and unfolded protein), cGMP-PKG, and cAMP signaling pathways (Fig. S7C). Most of these functions are related to stress response, indicating that the whole brain cells in our set environment are collectively involved in stress regulation after death. However, genes involved in synapses, neural development, and cell adhesion were under-expressed in most cells (Fig. S7D).

The expression changes of these co-DEGs in different PMIs were subsequently analyzed. We observed that the expression of these co-DEGs was not significantly increased or insufficient between H48 and H54 (Fig. S7E, F). Although the organism has been in the death state of ischemia and hypoxia, the gene expression activity continues and does not stop with the death of life.In addition, the expression of co-upDEGs was the lowest in Oligo at H0, but reached a comparable level as other cells at H24 (Fig. S7G), while the change of co-downDEGs appeared to be independent of cell type (Fig. S7H). We speculate whether it is due to the fact that the expression change of co-upDEGs is the greatest in Oligo leading to the most damaged Oligo cells. To test this hypothesis, fold change (log2FC) of gene expression between H24-H54 and H0 was calculated. Similar to the mentioned above, significant changes in co-DEGs occurred within 36h after death (Fig. S7I, J). Importantly, we found the most prominent changes in co-upDEGs in Oligo and OPCs (Fig. S7K), followed by Ast and Inh cells, which not only scored high in Augur analysis (Fig. S5), and the proportion change correlation of Inh cells was also second only to Oligo and OPCs (Fig. 2D). These results not only preliminarily validated our conjecture, but also confirmed the accuracy of the previous Augur analysis.

### 2.5 Oligodendrocytes and OPCs exhibit a PMI-dependent reactive signature

To identify transcriptional changes caused by PMI, we first analyzed oligodendrocytes and OPCs, which were preferentially effect. The more DEGs numbers of Oligo cells were observed with the extension of PMI, and the main contribution was down-regulated genes (Fig. S8A). Similar analysis of OPCs were performed, and the increased trend of down-regulated DEGs was consistent with Oligo, but more prominent (Fig. S8B), and nearly 45% and 80% of genes were co-up-regulated and co-down-regulated in all samples, respectively (Fig. S8C, D). Similarly, most of marker genes also revealed a trend of insufficient expression with the extension of PMI (S8E, F). We observed a qualitative change of these DEGs in H24 samples, and reached a plateau in H36, with almost complete up-regulation or down-regulation (Fig. S8E, F), indicating that partial genes have activities at the molecular level and play an unthinkable role under the condition of almost complete apoptosis of brain cells. Probably the low expression of most genes results in the inability of the Oligo cells and OPCs to sustain life operation, resulting in the cell death.

PMI-dependent Oligo cells-specific DEGs, including ribosomal genes (*Rps28*, *Rps27a*, *Rps24*, et. al.), various stress genes (*Cryab*, *Ubc*, et. al.) and long non-coding RNA (lncRNA) (*Snhg11*, *BC1*, *Tpt1*, *e*t. al.) were most upregulated in H24-H54 compared with H0 mice, and most of them are related to cerebral ischemia or hypoxia [25, 26]. The lncRNA *Snhg11* gene was highly expressed in the ischemic stroke brain [27], and *Tpt1* was a death associated promoter [28]. The downregulated genes, including *Pacrg*, *Unc5c* and *Xrcc6*, involved in regulating cell death (Fig. S9A). Such as, increased levels of *Pacrg* resulted in an increase in aggresome formation, and conferred significant resistance to aggresome disruption and cell death [29]; The Ku70 protein, a product of the *Xrcc6* gene, was recently identified as a critical anti-apoptotic protein [30, 31]. Thus, these specifically down-regulated genes apparently lose their resistance to cell death, resulting in preferential death of Oligo cells. Eigengenes of each group designated key gene expression trends, with subsequent Gene Ontology (GO) and KEGG analyses highlighting relevant biological pathways, The 68 co-up-DEGs in Oligo cells were mainly enriched in biological processes and pathways such as p53-mediated positive regulation of signal transduction, ribosomal subunit, microtubule cytoskeleton organization, stress response, cell necroptosis, axonogenesis and myelination (Fig. S8G). The 141 co-down-DEGs in Oligo cells were associated with Oligo cells mainly enclose axons in the central nervous system, form insulating myelin structures, assist in the efficient transfer of bioelectrical signals, and maintain and protect the normal function of neurons (Fig. S8H). The lack of oligo cells affects the signal transmission of neurons, so the gene expression of this pathway was insufficient. It has also been reported that glutamate excessively activates glutamate receptors on Oligo cells under pathological conditions such as hypoxic brain damage in brain tissue, leading to damage and death of Oligo and OPCs.

Likewise, the most significantly up-regulated DEGs in PMI-dependent OPCs were similar to those in Oligos, which may explain why these two cell types were most affected by PMI. However, the volcano plot of H24-54 versus H0 revealed a significant downregulation of three genes in particular- the cell adhesion (*Ctnna3*, *Ctnna2*, *Ctnna4*), protein autophosphorylation (*Fgfr2*) and synapse-related genes (*Mdga2*, *Grid2*), and the down-regulation was more significant in response to the prolongation of PMI (Fig. S9B). We then performed GO and KEGG analyses of co-up-regulated and co-down-regulated DEGs in OPCs. In addition to stress function, the co-upregulated DEGs of OPCs were also enriched in neuronal synapse-related functions and pathways (Fig. S8I). The co-down- regulated genes of OPCs were enriched in the functions of nervous system development, axon guidance, chemical synapses, cell adhesion and oligodendrocyte differentiation (Fig. S8J), suggesting that the differentiation ability of OPCs was weakened. Although the correlation priority of OPCs and PMI was not prominent, the cell loss was severe, which might be related to the differentiation capacity of OPCs themselves, and we suspect that they might play a role in recharge after brain tissue death. Although co-DEGs were removed prior to analysis of PMI-specific corresponding cells, the majority of up-regulated DEGs remained stress-related (ribosome function) genes in Oligo and OPCs compared with H0. We speculate that these cells are preferentially activated or respond to the adverse environment of postmortem ischemia.

To dissect the subclusters with the strongest response to PMI, taking advantage of the single- nuclei resolution of our data, we re-clustered Oligo cells to distinguish four sub-clusters (Fig. 3A-D), and found a cliff-like decrease in Oligo cell number with prolonged PMI, especially after 36h postmortem (Fig. 3E). We expected to identify the PMI-dependent Oligo subcluster, but all sub-cells showed similar death patterns and were almost synchronized (Fig. 3F, G). Furthermore, we interleaved marker genes of Oligo subpopulations and Oligo-DEGs to screen out the Oligo subcells- specific DEGs, and found that the Oligo-DEGs were also highly expressed in sub-cells (Fig. 3H), indicating that the differential expression of these genes and their functional changes caused by PMI promoted the death of Oligo cells. The expression of *Cd81* is increased in glial cells after hypoxic injury [32]. High expression of *Dst*, a neuroaxonal lesion marker, indicates that the destruction of Oligo cells leads to a decrease in the ability of myelination [33]. C-type natriuretic peptide (*Cnp*) plays a neuroprotective role after hypoxic-ischemic brain damage [34], and this gene is highly expressed in Oligo cells at 24h after death, and its expression increases with the increase of PMI. Perhaps this protective mechanism is not direct. However, it continues to act during hypoxia-ischemia. However, the expressions of *Mal* and *Cldn11* [35], which promote the formation of radial components of myelin sheath, are highly expressed in the hippocampus after death for a long time, and have a protective effect on neurons.

**Figure 3.**
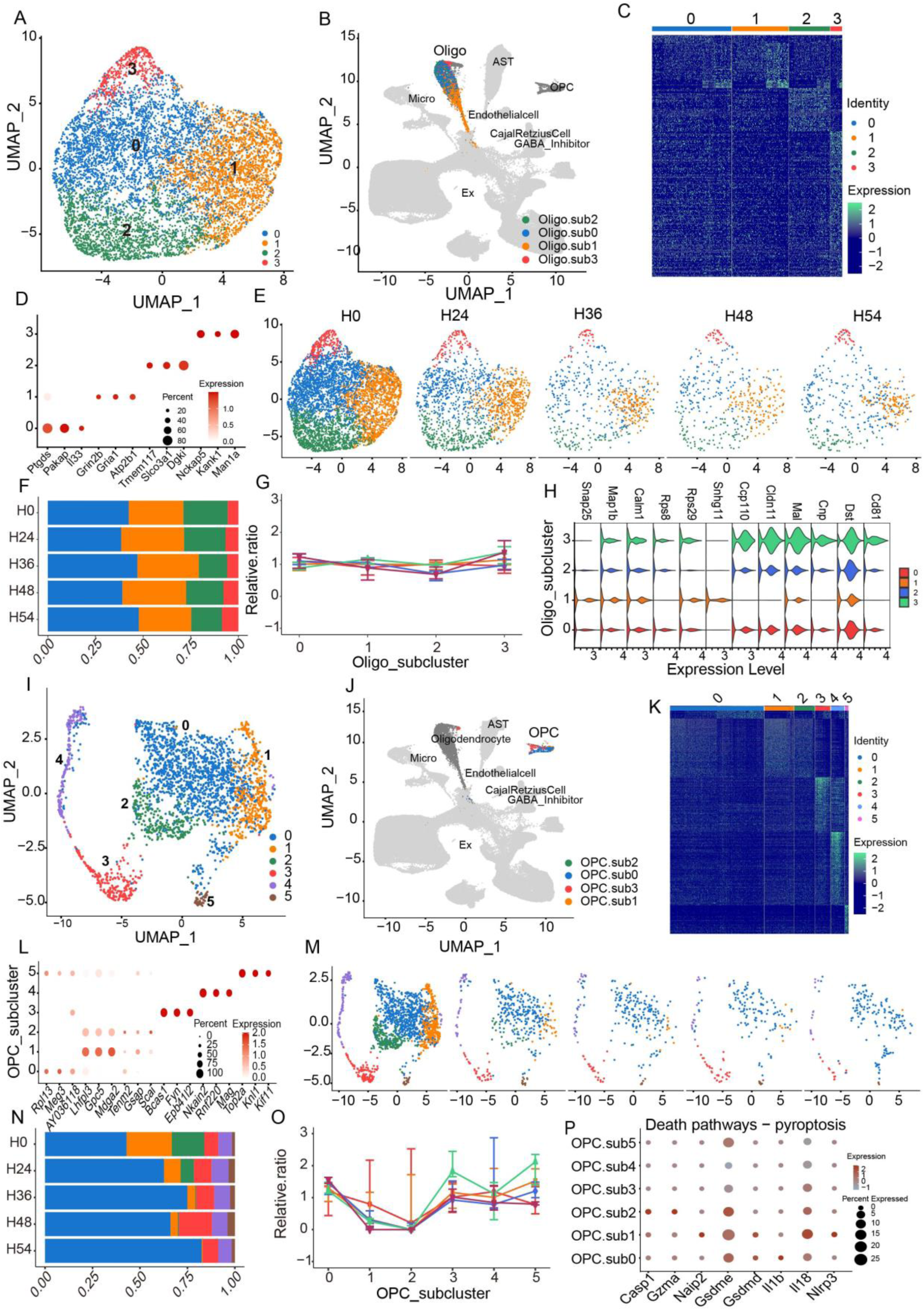
Characterization of the oligodendrocyte and OPCs subclusters. (A) UMAP of re-clustered oligo identifying 4 sub-clusters. (B) UMAP showing the oligodendrocyte cluster (cluster 4) from Fig. 1C. (C) Heat map showing the average gene expression of top DEGs (top (fold change) in the Oligo subclusters. Color scheme shows row max and row min, which represents relative expression of each gene among all samples. (D) Marker genes of Oligo subclusters states. Expression level (color scale) of marker genes across clusters and the percentage of cells expressing them (dot size). (E) UMAP across four major Oligo states in five PMI point. (G) Relative frequency of Oligo subclusters in F. Data are presented as mean ± s.e.m. (H) Violin plots showing the expression of PMI- dependent Oligo genes. Oligo DEGs were highly expressed in all its sub-cells. Violin plots are presented with floating boxes showing median (middle line) and quartiles (top and bottom). Minima and maxima are shown as the bottom and top of the violin plots. (I) UMAP of re-clustered oligo identifying 6 sub-clusters. (J) UMAP showing the OPCs cluster (cluster 11) from Fig. 1C. (K) Heatmap of average gene expression of top DEGs in the OPCs subclusters. (L) Marker genes of all OPCs subclusters. (M) UMAP across OPCs subclusters in five groups. (N, O) Frequency and relative frequency of OPCs subclusters. (P) Expression of pyroptosis pathway-related genes in OPCs subclusters, pro-inflammatory signaling triggered pyroptosis pathway were highly expressed in OPCs subcluter 1 and 2. n = 3 biologically independent mouse brain samples per group.

Next, we re-clustered OPCs and unsupervised differentiated six subclusters (Fig. 3I-L), and subclusters 1 and 2 (*Lhfp13*^+^_OPCs) belonged to the same genotype cells with most responsive to PMI, which were important contributors to the reduction of OPCs (Fig. 3M, 3O). Moreover, the specifically expressed genes of *Lhfp13^+^*_OPCs were associated with inflammation. The expression level of *Lhfp13* was increased in malignant glioma tissues [36], and glypican 5 (*Gpc5*) was a susceptibility gene for inflammatory demyelinating diseases [37]. The *Tenm2* gene has been reported to be associated with inflammation, and would induce microglia activation, to induce microglial activation in AD mice, and increased expression of AD-related genes of Oligo was partially dependent on *Trem2* [21]. In addition, we found that all the genes involved in the pro-inflammatory signaling triggered pyroptosis pathway were highly expressed in *Lhfp13^+^*_OPCs cells than other subpopulations (Fig. 3P), indicating that the loss of these two contributing clusters was due to inflammatory pyroptosis.

In summary, Oligo and OPCs were severely affected by postmortem time or sample degradation and preferentially died in brain tissue. More stress genes were highly expressed in these cells, indicating greater sensitivity to external environmental perturbations. Oligo and OPCs were also susceptible cells of hypoxic-ischemic stroke research, and we speculated whether the highly expressed genes of these cells in our data can also be used as candidate genes related to stroke disease.

### 2.6 Neurons show minimal transcriptional changes

We next examined neuronal and other glial cells, and found that cell types other than microglia and oligodendrocytes evinced more limited transcriptional responses to PMI. Neuronal cells account for the largest proportion of the brain, and are most significantly affected by hypoxia-ischemia in previous reports [24]. snRNA-seq data showed that the expression of most marker genes of Ex was relatively constant during postmortem (Fig. S10A). Compared with H0, the up-regulated DEGs were predominant, especially in H36 vs. H0 (Fig. S10C). The top up-DEGs were inconsistent with Oligo and OPCs, and were not stress-related genes (Fig. S10B). *Dynll1*, *Hpca*, *Ncdn*, *Pcp4* and *Calm3* were notably upregulated in H24-H54 samples compared with H0 (Fig. S10B). These up-regulated genes were mainly involved in neuronal projection development, synaptic vesicle endocytosis, glycolysis, ATP metabolism and ion transport processes. Calcium signaling, regulation of the actin cytoskeleton, mTOR signaling, Hippo signaling, and Apelin signaling were also enriched (Fig. S10E). The Hippo signaling pathway can regulate organs by regulating cell proliferation, apoptosis, and stem cell self-renewal ability, and we hypothesized that activation of these signals may protect nerve cells for a long time in vitro.

Then, we re-clustered Ex cells into 13 subclusters (Ex_sub.0-12) (Fig. S10F), and each Ex subcells was visually distributed equally across all groups, except for Ex_sub.5 and Ex_sub.7, which showed some sparsity (Fig. S10G). The detected frequency of most Ex subclusters was constant, but decreased in Ex_sub.5 and 7 clusters (Fig. S10H). However, the functions and death-related pathways specifically enriched in Ex_sub.5 and Ex_sub.7 were not fundamentally different from other sub-clusters (Fig. S11). In addition, we observed that the proportion of Ex_sub.1 increased with the prolongation of PMI and was more prominent in the H36 (Fig. S10H). This cluster accounted for a large proportion, and we speculated that the cell state change of Ex_sub.1 might be disturbed the percent of Ex_sub.5 and Ex_sub.7. As we suspected, Ex_sub.1 was specifically and independently enriched in functions and pathways related to stress-response (ribosome, thermogenesis, oxidative phosphorylation) and ATP metabolism terms (proton-driven ATP synthesis, proton transmembrane transport, ATP metabolic process, proton-driven mitochondrial ATP synthesis, and ATP biosynthesis process) (Fig. S10I). Comparisons with the Ex_sub.1 and Ex_sub.10, the other subcells revealed significant enrichment of gene sets nervous system development, ion transmembrane transport, axon guidance, cholinergic/glutamatergic synapse, cell adhesion and genes involved in the regulation of cell growth, differentiation, survival and stress response pathways (Fig. S11). The specific DEGs in Ex_sub.1 mainly included eukaryotic translation elongation factor (*Eef* ^+^), ribosomal protein (*Rps*/*Rpl* ^+^) and ATP related (*Atp* ^+^). Except for ATP-related genes, the remaining DEGs of Ex_sub.1 showed a higher similarity to the top DEGs of Oligo and OPCs. Meanwhile, Ex_sub.10 and Ex_sub.1 showed similar GO and KEGG terms enrichment (Fig. S11), and were the subpopulations with the largest response to PMI. In addition, endogenous apoptosis, ferroptosis, and necroptosis pathways related genes were higher expression in Ex_sub.1 and Ex_sub.10 (Fig. S10J), these cells should be preferentially lost based on the pattern of changes in Oligo and OPCs, but inconsistent with our results. We hypothesized that Ex_sub.1 might preferentially perceive the changing environment of the brain, then take stress measures, and drive ATP to provide more energy to maintain the life characteristics of cells. The Ex_sub.1 increased temporarily in H24 and H36, but the subsequent energy supply may continue to be insufficient, and loss the ability to compete with the outside world leads to the fluctuation of the cell proportion. This might be the reason why cell state changes of Ex_sub.1 were inconsistent with OPCs and Oligo.

Subsequently, similarly analysis was unfolded for AST, Micro, Inh, and Endo cells. AST with the high augur prioritization score, and AST_sub.2 was the cluster with the most prominent state change (Fig. S12A-D), which significantly upregulated genes of ionotropic glutamate receptor signaling pathway (Fig. S12E). Moreover, the effect of PMI on different AST subcells was poles apart (Fig. S12E), and was the only cells that was so affected. Cell adhesion and junction related pathways enriched in AST_sub.1, ribosome and positive regulation of signal transduction by p53 class mediator enriched in AST_sub.3, cell differentiation and development, glutamatergic and cholinergic synapse related pathways enriched in AST_sub.4 (Fig. S12E). We also observed that AST_sub.3 was distributed independently from other subclusters, and the top DEGs of H24-54 vs. H0 from AST were more and highest expressed in AST_sub.3 (Fig. S12F). *Gnas* will activate adenylyl cyclase in the G protein-coupled receptor signaling pathway, leading to an increase in cAMP levels and participating in the regulation of cell growth and cell division. Downregulation of *Gnas* in vitro would reduce cell apoptosis [38], and up regulation of *Gnas* was confirmed to accelerate cell death activity in substantia nigra of Parkinson’s disease [39]. *Olfm1* has been reported to be associated with cell viability, and was confirmed to induce cell necrosis and apoptosis [40]. More ribosomal genes were highly expressed in AST_subcluster 3, similar to the PMI-dependent subclusters of oligo and OPCs. Subsequently, similar subtype analysis was performed for the remaining cell types, in which the most disturbed cell subclusters may be derived from the differences caused by sample 1, such as Micro_sub2 and 3 and Endo_sub2 and 5 (Fig. S13). Overall, these cell types were less affected by PMI, and a significant proportion of cell subtypes were detected at all postmortem time points.

### 2.7 Ribosomal gene expression changes dependent on PMI

In the process of extensive manual screening of data, we discovered a gene set - ribosomal protein genes (*Rpl* and *Rps*, RB genes), which were easily overlooked in conventional analysis. These genes were significantly up-regulated in most cell types (Fig. S6 and S7), and we proposed a bold hypothesis - perhaps the expression changes of RB genes are related to cell fate? Perhaps one of the key factors explaining the different sensitivity of diverse cell types to the PMI. We screened out the RB genes from all datasets and found that their expression changes over PMI were consistent with co-DEGs, that is, the expression of RB genes upregulated with PMI and terminated at H48 samples (Fig. 4A). The expression of RB genes was then analyzed in all cell types, and very low at 0h postmortem, except Micro and Endo, while gradually increased with the prolongation of PMI, especially from 24h after postmortem (Fig. 4B). Among them, the expression of RB genes was lowest in Oligo and OPCs at H0, but was comparable with other cells at H24 and reached a maximum at H48 (Fig. 4C). The function terms of these RB genes enrichment were almost consistent with co-upDEGs, including translation, ribosome and ubiquitin ligase related functions (Fig. 4D). Subsequently, we observed no significant increase in the expression of RB genes at H48 compared with H36, implying that the significant changes in these genes, although terminating at 48h, both reached a plateau at 36h (Fig. 4E). As expected, the log2(FC) of RB gene expression was highest in Oligo and OPCs, followed in AST and Inh cells (Fig. 4F), consistent with Augur scoring (Fig. S5A). In addition, the numbers and average log2(FC) of up-regulated RB genes, heat shock protein genes, and ATP-related genes in all cell types were calculated. The RB genes and ATP-related genes were upregulated most in neurons, but changed most prominent in Oligo and OPCs cells. The heat shock protein genes for the stress response changed similarly in all cells except neurons (Fig. 4G, H). We speculate that all glial cells receive stress signals first in the postmortem ischemic state, preferentially protecting the normal information transmission of neuronal cells. In short, RB genes showed the greatest changes in Oligo and OPCs cells, which seems to tentatively validate our previous hypothesis.

**Figure 4.**
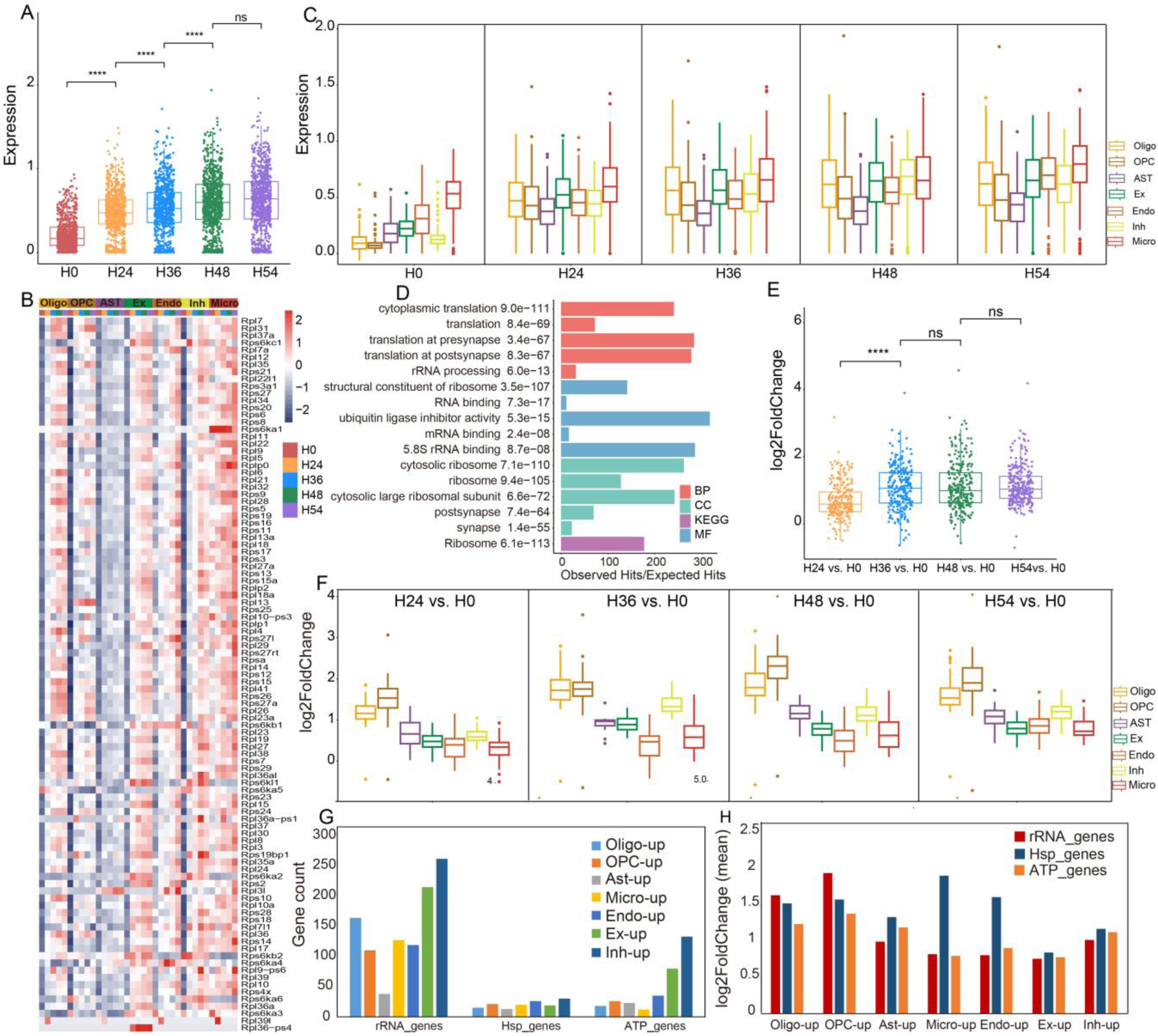
Expression analysis of ribosomal genes in all cell types. (A, B) Scatter plot and heatmap of all *Rps*^+^ and *Rpl*^+^ gene expression in five PMI samples (A) and cell types (B). (C) Boxplot of *Rps*^+^ and *Rpl*^+^ expression in all cell types of five groups. (D) Functions and pathways enriched by *Rps*^+^ and *Rpl*^+^ genes. (E, F) Log2(FC) of *Rps*^+^ and *Rpl*^+^ genes expression in all groups (E) and cell types (F). (G, H) The number (G) and Log2(FC) (H) of ribosomal genes, heat shock protein genes and ATP-related genes in all cell types, respectively. Log2(FC), log2 fold change. ns, not significant (*p* >0.05).

### 2.8 Variation of ribosomal genes in cellular subpopulations

To further confirm our hypothesis, we immediately analyzed the expression status of ribosomal genes in various cell subclusters, and we first focused on oligo and OPCs. The expression of RB genes in Oligo cells gradually increased with the prolongation of PMI and was similar in each of the subclusters of five group samples (Fig. 5A, B). The results of change fold imply that the expression change of RB gene reached a plateau at 36 h, which was consistent with the previous conclusion (Fig. 5C, 4A). The similar log2(FC) implies that the RB gene exerts a nondiscriminatory effect on the cellular state of the Oligo (Fig. 5D). We then performed a similar analysis on OPCs, and the expression of RB gene also increased with the increase of PMI in OPCs subcluster cells (Fig. 5E), but still did not reach the plateau after 36 h, and continued to increase significantly (Fig. 5G). Simultaneously, the most preferentially changes of OPCs_sub1 and OPCs_sub2 were observed in H24, and the change of OPCs_sub1 remained ahead in the subsequent sample group until the cells were undetectable, but could not be observed in the later comparison due to the absence of OPCs_sub2 in H36 (Fig. 5H). Excitingly, this was consistent with the subcluster proportion of OPCs, where OPCs_sub1 and 2 present a state of cell loss at different postmortem times (Fig. 3N, O). Subsequently, the same analysis strategy to examined the remaining cell subclusters. Among them, the largest fold change in RB gene expression was observed in Ex_sub5, Ex_sub7 and Ex_sub10 of excitatory neurons, Inh_sub2 of inhibitory neurons, Ast_sub3 of astrocytes, Micro_sub6 and Micro_sub7 of microglia, and Endo_0 and Endo_5 of endothelial cells (Fig. S14). In contrast to H0, most of these cells were missing in postmortem brain tissues, which would again confirm our assumption.

**Figure 5.**
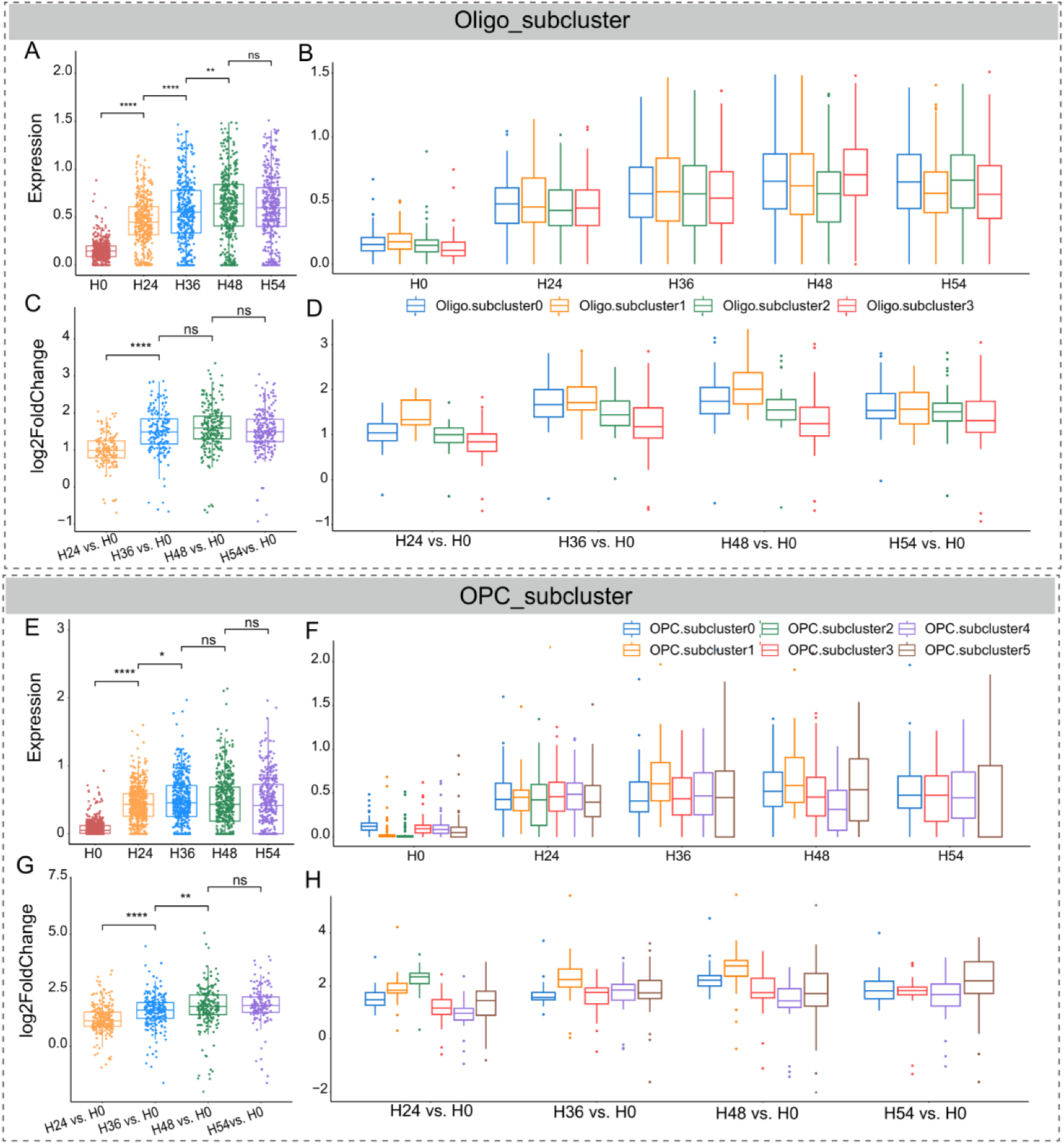
Differential expression analysis of ribosomal genes in the subpopulations of oligodendrocytes and OPCs. (A, B) ribosomal gene expression in Oligo and its sub-cells at 5 time points. (C, D) Log2(FC) analysis of ribosomal genes in A and B. (E, F) ribosomal gene expression in OPCs and its sub-cells. (G, H) Log2(FC) analysis of ribosomal genes in E and F.

### 2.9 Histological and cell death processes analysis

To confirm the damage of PMI to the nucleus, we first investigated cytoarchitectural changes of hippocampus using unbiased sampling and histological analysis. We analyzed brain cytoarchitecture using H&E and Nissl staining in the hippocampus–a region that are particularly susceptible to ischemia [8, 9]. The cell membrane of three regions was disrupted and the nucleus showed pyknosis in H24 to H54 group (Fig. S15A-C), indicating the existence of an apoptotic process. In addition, cells may reduce RNA synthesis and other nuclear activities when they are hypoxia, and we observed smaller nuclei. However, CA1 and dentate gyrus (DG) nuclei counts did not diminish as ischaemic durations increased, only CA3 nuclei numbers decreased significantly (*p* < 0.05) at H24, and with comparable depletion in H24 to H54 groups (Fig. S15D-F). Mammalian cells require oxygen to maintain cellular and tissue viability [41]. In just minutes after ischaemia, intracellular acidosis and oedema develop and trigger secondary injury to membranes and organelles, often causing cell death [24]. However, compared with brain cells, long-term postmortem ischemia has less damage to the nucleus and has regional differences. As an indicator of early cytotoxic oedema from ischaemic stress, ellipsoid cell morphologies increased with longer PMI in the CA3 (Fig. S15I), but experienced a bottoming effect at all time points in the CA1 and DG (Fig. S15G, H), possibly stemming from too long postmortem intervals. Moreover, Nissl bodies, which are essential components of protein synthesis in neurons, are significantly decreased when neurons are stimulated. The result of Nissl staining indicates that the nuclei numbers of neuronal was less affected by the PMI, but the cells in the CA1 and CA3 were more damaged with the prolongation of PMI, and the DG cells appeared to be more vulnerable to damage (Fig. S15G-I). This indicates that PMI has a greater impact on neuronal cells, which is consistent with previous reports [24], but may have less impact on the nucleus.

We next investigated key proteins of cell death pathways using immunohistochemistry analysis. For apoptosis, we measured the immunolabelling intensity of activated caspase-3 (act-CASP3) and performed a TUNEL assay. The analysis of the intensity of act-CASP3 immunolabelling of CA1 and DG was lower than in CA3 at H0 group, and observed their significantly increased (*p* < 0.05) as ischaemic durations up to H48 (Fig. S16A, F, G, L and Fig. S17A, F). The postmortem ischemic state activated caspase-3, and we suspect that the apoptotic process occurs almost throughout the experimental setup. It has been reported that caspase-3 was difficult to detect in tissues beyond 8 h postmortem, but our assay negates this statement. Conversely, the intensity of the TUNEL assay had a nearly to the levels trend in H24 and H0, and increased progressively in the other groups (Fig. S18). In addition, we were curious whether there was a pyroptosis process (a cell death pathway triggered by proinflammatory signals) during PMI, and we performed interleukin-1β (IL-1B) immunohistochemistry analysis. Across the investigated regions, IL-1B immunolabelling intensity was comparable in the H24 and H0 of CA1, and increased to a peak at H36 in CA1 and DG and subsequently stabilized or decreased, whereas the increase in intensity was more persistent in cells in CA3 (Fig. S16B, F, H, L and Fig. S17B, F). Subsequently, we investigated necroptosis, ferroptosis and autophagy, three distinct cell death pathways, using immunohistochemistry analysis of the important proteins in these pathways-receptor-interacting serine/threonine kinase 3 (RIPK3), the glutathione peroxidase 4 (GPX4) and beclin-1, respectively. The results were similar between the three cell death pathways and among all of the regions, but ferroptosis pathway seems to occur later, and GPX4 is stable within 36h after death (Fig. S16D and Fig. S17C). Compared with the H0 group, immunolabelling intensity was significantly increased in the other groups. In general, cells of hippocampus underwent various death processes during PMI, and most of them peaked at H36, and the susceptibility of the three regions to different death processes was different (Fig. S16 and S17). However, surprisingly, the effect on the total number of nuclei and neuronal nuclei was weak, which may also be the factor for the success of snRNA-seq.

## 3 Discussion

The sudden removal of brain tissue in vivo is in many ways mimics a catastrophic event that occurs with a hypoxic brain injury or a traumatic death with exsanguination. The energy demands of brain tissue are high, estimated to be ten times higher than those of other tissues, and the effect of energy deprivation on single molecules due to the postmortem time after the tissue is isolated is unknown. In this study, as a means to understand how PMI selectively affects cellular composition and gene expression, we performed snRNA-seq and histological analysis of mouse hippocampal tissues at 25°C with different PMI.

To isolate RNA that accurately reflects the transcriptome at the point of harvest, raw biological samples should be processed by freezing in liquid nitrogen, immersing in RNA stabilization reagent or lysing and homogenizing in RNA lysis buffer containing guanidine thiocyanate as soon as possible [42]. However, in certain environments or conditions, complete protective measures cannot be implemented in isolated tissues, resulting in RNA degradation and transcriptional differences. The RNA integrity of biological tissues is an important factor affecting the results of transcriptome sequencing data. However, there is still a lack of transcriptomic data on RNA degradation caused by postmortem ischemia time at the single-molecule level. In our study, snRNA-seq was used for the first time to comprehensively investigate the effects of PMI on cellular composition and gene expression at single-molecule resolution in the mouse postmortem brain hippocampus. The time lag between sample harvest and RNA protection influences expression of specific genes. Thus, when setting up the control and experimental groups, we strictly controlled the experimental process and conditions, ensured the consistency of sample processing methods, and marked the accurate sampling and storage time of each sample. In addition, the susceptibility of the brain to ischaemic injury dramatically limits its viability following interruptions in blood flow. Complete cerebral ischaemia paradigms have established that brain function ceases within seconds of cerebrocirculatory arrest, with high-energy metabolites depleted within minutes [43]. Even 5 min of global cerebral ischaemia can induce delayed neuronal death in certain selectively vulnerable brain regions. Furthermore, the susceptibility of the brain to ischaemic injury dramatically limits its viability following interruptions in blood flow. Complete cerebral ischaemia paradigms have established that brain function ceases within seconds of cerebrocirculatory arrest, with high-energy metabolites depleted within minutes [43]. Even 5 min of global cerebral ischaemia can induce delayed neuronal death in certain selectively vulnerable brain regions. The pathophysiological mechanisms of global cerebral ischemia mostly come from animal models constructed by various methods. Studies on the neck amputation model have confirmed that the brain has a highly limited energy reserve, which may be the basis of its susceptibility to ischemic or hypoxic conditions. However, the brain was harvested by decapitation in our study based on the constraints of the experimental conditions and the tradeoff of the advantages and disadvantages of each approach.

Previous studies have shown that cryopreservation at room temperature (warm and cold ischemia) affects the integrity of extracted RNA [44], and the storage times also significantly affect RNA quality and gene expression levels [14]. Auer et al. showed that the storage temperature has a significant effect on RNA yield and integrity [45]. In addition, several studies have reported that the use of RNA samples with RIN values of around 5.0-7.0 does not negatively affect estimates of the gene expression profile [46] or *de novo* assembly [47]. However, some results indicated that even if using high-quality RNA (RIN > 9.1) for RNA-Seq, the gene expression levels of specific genes were significantly different between samples that share the same cultivation day but have different processing lag times [42]. In our data, RIN dropped to about 3 at the postmortem interval of 54h, which is clearly risky to adopt samples with RNA fragmentation, and there was also concern before our experiments, that samples with low RIN values can be able to perform snRNA-seq smoothly? However, the exciting results of cDNA fragment distribution disproved the inherent understanding, and also broke through the shackling of high-throughput snRNA-seq platform on sample RIN limitation. The postmortem brain tissues are stored at room temperature, and the mRNA is more constant than the rRNA within nuclei, which is contrary to our previous effect of formaldehyde fixation time on these RNAs [19]. It also indicates that RIN can only be used as an indicator of RNA quality in ideal archived tissues for basic experiments, and its guidance for transcriptome research of normal temperature or fixed samples are limited. Taken together, our data for the first time demonstrated the presence of excellent transcriptional mRNAs in low RIN samples for successful snRNA-seq applications.

Our snRNA-seq data revealed a significant upregulation of RP genes in almost all cells, with the expression variation of RP gene set being more pronounced than that of other genes in response to PMI in brain tissue. In addition, the expression changes of RP gene-set was most significant in Oligo and OPCs, and was also observed in cell subsets with large cell state perturbations. It has been reported that ribosomal protein genes can provide a better starting point for understanding gene regulation and constructing gene regulatory networks [48], indicating the importance of ribosomal genes in living organisms. Moreover, mutations or abnormal expression of ribosomal genes have a wide range of effects on the physiological functions of cells, leading to a variety of serious consequences (cell metabolism time, cell growth arrest and even death, etc.). Aberrant expression of ribosomal genes has been reported to be associated with genetic diseases. In 1990, Fisher published a thesis in Cell that *Rps4* gene deletion was an important cause of Turner syndrome [49]. The mutation of *Rps17* gene plays a role in the occurrence of Diamond Blackfan anemia [50]. In addition, congenital blepharoptosis (*Rps8*), Bardet-Biedl syndrome 1 (*Rps30*), congenital fatal contracture syndrome (*Rps12*), Camurati-Engelmann disease (*Rps18*), dystrophic myotonia (*Rps32*), Bardet- Biedl Syndrome 4 (*Rpp1*) is associated with abnormalities in ribosomal genes [51]. Moreover, the abnormal expression of ribosomal genes plays an important role in the regulation of cell division, proliferation and differentiation, and also plays a role in the occurrence and development of tumors. For example, after malignant transformation of BEP2D cells, *Rpll0*, *Rpl31*, *Rps23* and *Rpl22* were highly expressed. Sudden removal of brain tissue from sudden death mice mimics in many ways the catastrophic events that occur due to anoxic brain damage or traumatic death due to blood loss, and the energy requirements of brain tissue are high, estimated to be 10 times higher than those of other tissues [52]. Studies have shown that the expression of 60 ribosomal protein genes in zebrafish gills under hypoxia stress is significantly up-regulated, and GO enriched pathways include translation, protein assembly and ribosome [53]. Therefore, we hypothesized that ribosomal genes may be one of the important factors leading to the change of cell state, and their overexpression may play a regulatory role in cell death.

In rodents, post-mortem slice cultures have been paired with electrophysiological interrogations to reveal that neuronal activity is recoverable even up to 6h after death [54]. In addition, viable tissue specimens of the developing human brain can be harvested even 18-49 h after death following cold storage for use in neural cell proliferation and differentiation studies [55, 56]. As observed in ex vivo cell/tissue culture and brain studies, along with the selective cerebrocirculatory arrest models, neural cells and tissue can withstand prolonged periods of ischaemia in the absence of intravascular blood. These studies indicate that brain cells and tissue can tolerate prolonged periods of ischaemia, retaining their capacity for cellular recovery under the appropriate circumstances. In our date, although the characteristics of Ex-sub1 and 10 were similar to those of Oligo and OPCs, the highly expressed genes in the remaining excitatory neural subsets were enriched in terms of Ca^2+^ binding, calmodulin binding, calcium signaling pathway, voltage-gated ion channel activity, ion channel activity and glutamatergic synapses. Ischaemia would increases cytosolic Ca^2+^ concentration, and many channels and pathways have been implicated in mediating the effects of supraphysio logical increases in Ca^2+^ concentration in various models of anoxia and ischaemia, including ionotropic glutamate receptors and voltage-gated calcium channels. Increased cytosolic Ca^2+^ concentration during ischaemia also drives the release of excessive glutamate, and the mechanisms underlying glutamate-mediated Ca^2+^ excitotoxicity comprise a wide range of cellular processes, including generation of ROS/RNS, activation of lipases and proteases, mitochondrial dysfunction and apoptosis [57]. Although our data did not show a strong perturbation of the state of excitatory neurons, apoptosis is inevitable. Astrocytes was reported less sensitive than neurons to hypoxic conditions in vitro [58], possibly owing to the former’s ability to upregulate glycolytic capacity during hypoxia [59]. And during permanent focal ischaemia in rats, astrocytic decline may precede neuronal demise within the lesion core [60]. However, our Augur analysis showed that the dependence of AST on PMI was almost weaker than that of Oligo. Therefore, the priority and mechanism of the effects of postmortem ischemia and hypoxia on neurons and AST cells need to be further explored. Microglia, the resident immune cell of the brain, under physiological conditions, microglia usually exhibit a “resting” state, which is consistent with our results. Micro may sense local microcirculatory dynamics, and potentially halt rearrangements of their cytoskeleton to conserve ATP for subsequent activation upon reperfusion. In vitro, human umbilical vein endothelial cells are relatively resistant to hypoxia, with more than 90% of these cells exhibiting signs of viability after 6h of reduced oxygen levels, and approximately 50% of cells remaining viable after 24 h of hypoxia [61]. In addition, ischaemia disrupts normal endothelial production of nitric oxide (NO), a major physiological vasodilator, thereby exacerbating vascular dysfunction. Endothelial NO synthase (NOS) is the primary source of endothelial NO, and its biochemical function depends on the presence of molecular oxygen. Although under anoxia NOS can produce NO by reducing nitrite, it can do this for only a limited time. Dynein light chain 1 (*Dynll1*) was overexpressed in excitatory neuronal cells, which physically interacts with neuronal nitric oxide synthase, and we speculate that this binding may supplement NO deficiency due to hypoxia, and thereby protect neurons.

It is well known that removal of brain tissues from their normal environment can lead to death of cells once blood circulation is no longer available to oxygenate the tissue. It has been reported that the trend of gene set down-regulation in human brain neurons increased with the extension of postmortem time, while a reciprocal and dramatic increase was observed in glial cells [18]. However, in our single-molecule data, the opposite result was observed. Neuronal cells were dominated by up- regulated genes, whereas glial cells were mostly under-expressed and enriched in adhesion-related functions, which were enriched for neuronal down-regulated genes in above reported. In addition, the expression changes of housekeeping genes were not significantly correlated with PMI, which is consistent with previous reports, but not for all housekeeping genes. For example, Albde gene expression peaked at 24h and then dropped sharply, while Actb and Gapdh reached the ceiling high expression effect at H24. The time to reduce the body to 4°C during PMI is an important measure of RNA stability and a major factor in proteome degradation [62]. To date, no validated and accurate model of postmortem brain temperature is available, but a decrease of 1°C per hour has been reported in the first 12 hours immediately after death, followed by a decrease of 0.5°C per hour [63]. Even if the human body is refrigerated, the brain takes 30 hours to cool down. Thus, while we examined the effects of simulating PMI at room temperature, in most cases postmortem brains will be exposed to higher temperatures for longer periods than we used in this study and thus may undergo faster changes. Although some studies have shown that the RNA transcriptome is stable up to 30 hours after death, the dynamic changes in RNA levels in specific cell types as a result of PMI described here have been reported by others, showing RNA degradation, chromatin modification, 6 activation of gene expression, and protein degradation. Although we did not directly observe protein expression, predicted cellular differences involving specific cell populations may produce corresponding proteomic changes in glial and neuronal cell populations.

In conclusion, the computational analysis of snRNA-seq data revealed major transcriptomically defined cell types that were comparable to publicly available mouse single-cell datasets. The diverse cells have a different response over the time elapsed since death, and oligodendrocytes and OPCs responded most strongly to PMI, and severe loss occurred almost at the earliest stage after death, but neurons, astrocytes and microglia were less affected. In addition, the genes most affected by PMI were ribosomal genes, and this gene set was more significantly upregulated in cell types with the greatest cell perturbation. This extensive data resource complements and expands on previous studies and enables the systematic investigation of transcriptomic changes in multiple cell types exposed to different PMIs.

## 4 Methods

### 4.1 Ethics Statement

The study was approved by the animal ethical and welfare committee of Zhongda Hospital Southeast University (20200104005). All procedures were conducted following the guidelines of the animal ethical and welfare committee of SEU. All applicable institutional and/or national guidelines for the care and use of animals were followed. In addition, we sought to minimize the animal number and any potential discomfort and suffering.

### 4.2 Animal anaesthesia and surgical protocol

Eight-week-old male C57BL/6J mice were purchased from the Qing Long Shan Dong Wu Fan Zhi Chang, Nanjing, China The animals were anesthetized with 500 mg/kg tribromoethanol (Sigma, Saint Louis, MO, USA) and were killed by cervical dislocation. After the animals were sacrificed, brain tissues (hippocampus) were isolated, placed in 2 mL enzyme-free centrifuge tubes, and stored at 25°C at constant temperature. Postmortem tissues were sampled at 0 h (snap frozen, H0), 24 h (H24), 36 h (H36), 48 h (H48), and 54 h (H54), followed by snap frozen in liquid nitrogen and stored at -80°C until use. For the reliability of the data, we maintained the consistency of the environment for each sample and set up biological replicates of three mice per group.

### 4.3 Sample quality control and nucleus preparation

RNA was extracted from trace samples at different postmortem intervals using the QIAGEN RNA Extraction kit (74204), and RIN measurements were performed using an Agilent 2100 Bioanalyzer System. We selected five sets of samples with full RIN (9-3) coverage for nucleus preparation. Nucleus was isolated using Shbio Nuclei Isolation Kit (SHBIO, #52009-10, China). Briefly, the frozen tissue samples were quickly added to the lysis buffer and the tissue was ground to liquid state using a tissue homogenizer. The tissue lysate was passed through a 40 μm cell sieve to remove impurities and transferred to a new 2mL EP tube and centrifuge at 500 g at 4 °C for 5 min to obtain the precipitation. Next, PB1, PB2, PB3 solutions were added to the precipitation and the nuclei are located at the junction of PB2 and PB3 solutions. Finally, the nuclei were resuspended by blowing into 50 μL NB solution. Nuclei were counted with a cell counter (Thermo Fisher, America).

### 4.4 Library Construction and Sequencing

Sorted nuclei were processed using the 10×Chromium Single Cell 3′ Library and Gel Bead Kit v3, nuclei were immediately loaded onto a Chromium Single Cell Processor for barcoding of RNA from single nuclei. The quality of cDNA was assessed using the Agilent× v3.1 to generate the cDNA libraries. The quality of cDNA was assessed using the Agilent 2100 Bioanalyzer System. Sequencing libraries were constructed according to the manufacturer’s instructions and then sequenced in a NovaSeq 6000 sequencing system (Illumina, America).

### 4.5 Bioinformatics analysis

#### Data Demultiplexing and Quality Control

**Raw sequencing data was processed by** Cell Ranger 5.0.1 (10 × Genomics), the function count was used to align reads to the reference genome (mm10) with default parameters, and the R package Seurat v4.0 (PMID: 34062119) was applied for downstream analysis. Before we started downstream analysis, we focused on four filtering metrics to guarantee the reliability of our data: (1) ambient RNA contamination was estimated and removed by SoupX pipeline in R (PMID: 33367645); (2) nuclei with less than 200 genes and genes detected in fewer than three nuclei were filtered to avoid cellular stochastic events; (3) nuclei with a percentage of expressed mitochondrial genes greater than 5% were removed to rule out apoptotic cells; (4) potential doublets were predicted and removed by R package DoubletFinder V2.0 (PMID: 30954475). After filtering cells and genes according to the metrics mentioned above, there were 22,431 genes and 123,464 nuclei left for downstream analysis.

#### Sample integration, dimensional reduction and clustering

Samples were normalized using a scaling factor of 10000, and the FindVariableFeatures function was performed to determine the set of highly variable genes (n = 2000). Samples were then integrated using FindIntegrationAnchors and IntegrateData functions in Seurat. ScaleData and RunPCA were performed and the top 22 principal components were used based on the ElbowPlot. Dimensional reduction and clustering were performed using the FindClusters function (resolution 0.1) and projected to two-dimensional uniform manifold approximation and projection (UMAP) embedding images.

#### Marker identification and annotation

The marker genes of each cell type were calculated through the FindAllMarkers function in Seurat, the threshold for log2 fold change (logFC) was 0.5 and the adjusted P value was 0.05. Then cell type was annotated based on the markers from the previous studies and the CellMarker 2.0 database (PMID: 36300619). Biological functions and pathways were analyzed by enrichment analysis using the Gene Ontology (GO) and Kyoto Encyclopedia of Genes and Genomes (KEGG) databases by clusterProfiler package (PMID: 22455463).

#### Statistical analysis

The statistical and bioinformatics analyses, data visualization and plotting were conducted using R software (version 4.3.2). The Wilcoxon signed-rank test was used to investigate differences between the two groups.

### 4.6 Histological data analysis and quantification

#### H&E and cresyl violet (Nissl) staining assessment of the nuclei numbers

All staining was done through Servicebio, Wuhan, China. All of the H&E and Nissl slides were scanned. Two images were taken per region of interest (hippocampal CA1, CA3 and DG regions) and were evaluated using the cell counter function in ImageJ.

#### Cell death pathway quantification and analysis (actCASP3, IL-1B, RIPK3, GPX4 and Beclin-1)

For the CA1, CA3 and DG regions, three images were taken randomly from each slide and from the corresponding areas. The actCASP3^+^, IL-1B^+^, RIPK3^+^, GPX4^+^ and Beclin-1^+^ cell intensity was quantified manually by a blinded observer using ImageJ.

#### TUNEL quantification and analysis

All of the images were taken using a fluorescence microscope followed by a random same-size snapshots of the slides. The intensity was quantified manually by a blinded observer using ImageJ.

## CRediT authorship contribution statement

Yunxia Guo: investigation, methodology, validation, writing-original draft. Hao Huang and Junjie Ma: Data Curation and visualization. Xiaoying Ma, Kaiqiang Ye, Chao Jiang, Jitao Xu, Yan Huang, Xi Yang, Qinyu Ge, Jianyou Zhang and Guangzhong Wang: Writing-review & editing. Xiangwei Zhao: Methodology, supervision, resources, project administration and funding acquisition. All authors had read and agreed to the published version of the manuscript.

## Data availability

The single nuclei RNA sequencing data that support the findings of this study have been deposited in the National Center for Biotechnology Information (NCBI).

## Acknowledgments

This study was supported by the National Natural Science Foundation of China (81827901, 82361138570 and 32371393).

## Supplementary Material

**Figure S1.**
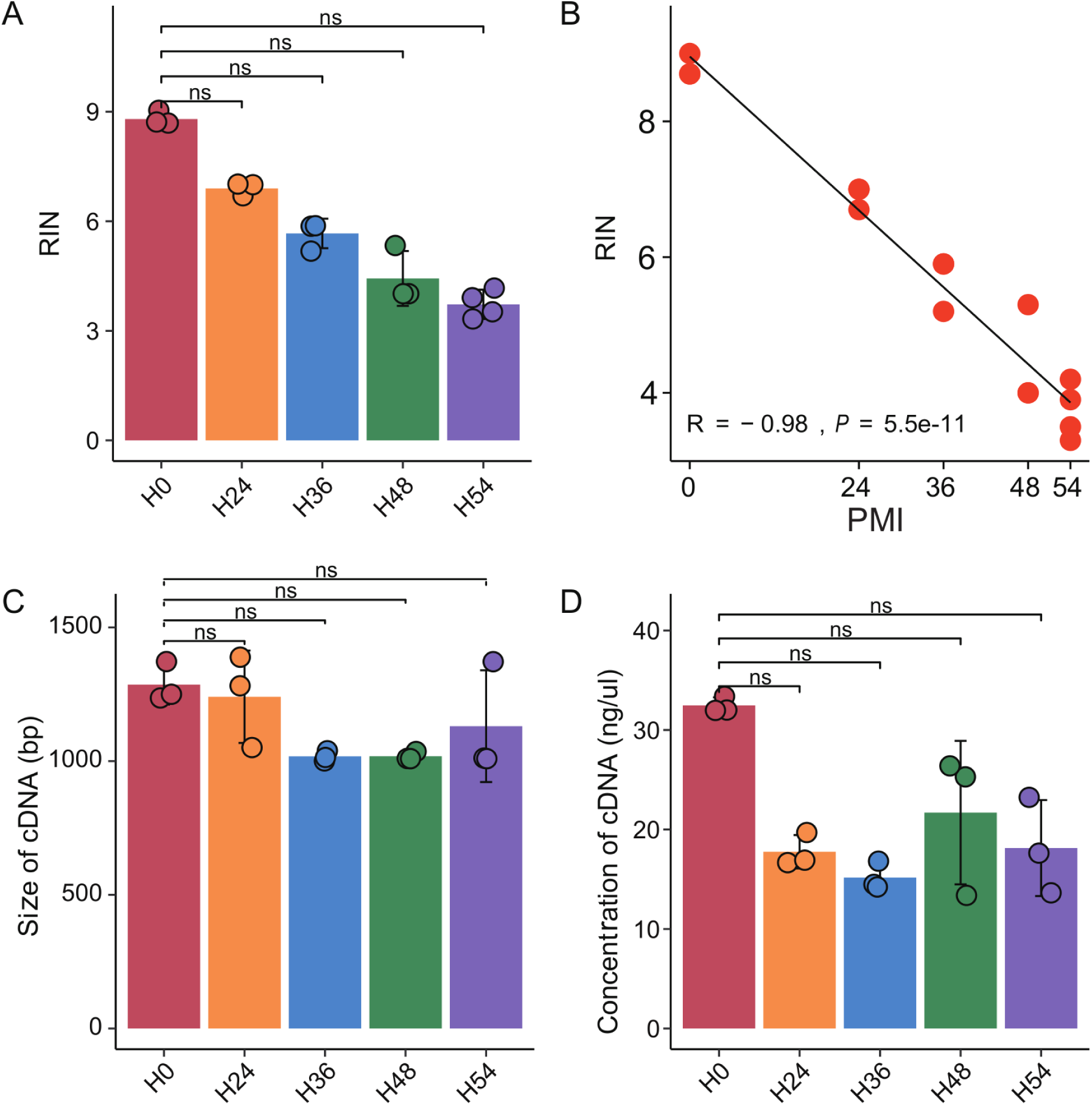
Effect of different postmortem cold ischemia times on RNA quality in brain tissue. (**A**) The effect of PMI on RNA integrity numbers (RIN). (**B**) The correlation between PMI and RIN values. (**C, D**) Representative peak values of amplified cDNA and its yield in five groups. n = 3 biologically independent mouse brain samples per group. Bars show mean ± s.d.

**Figure S2.**
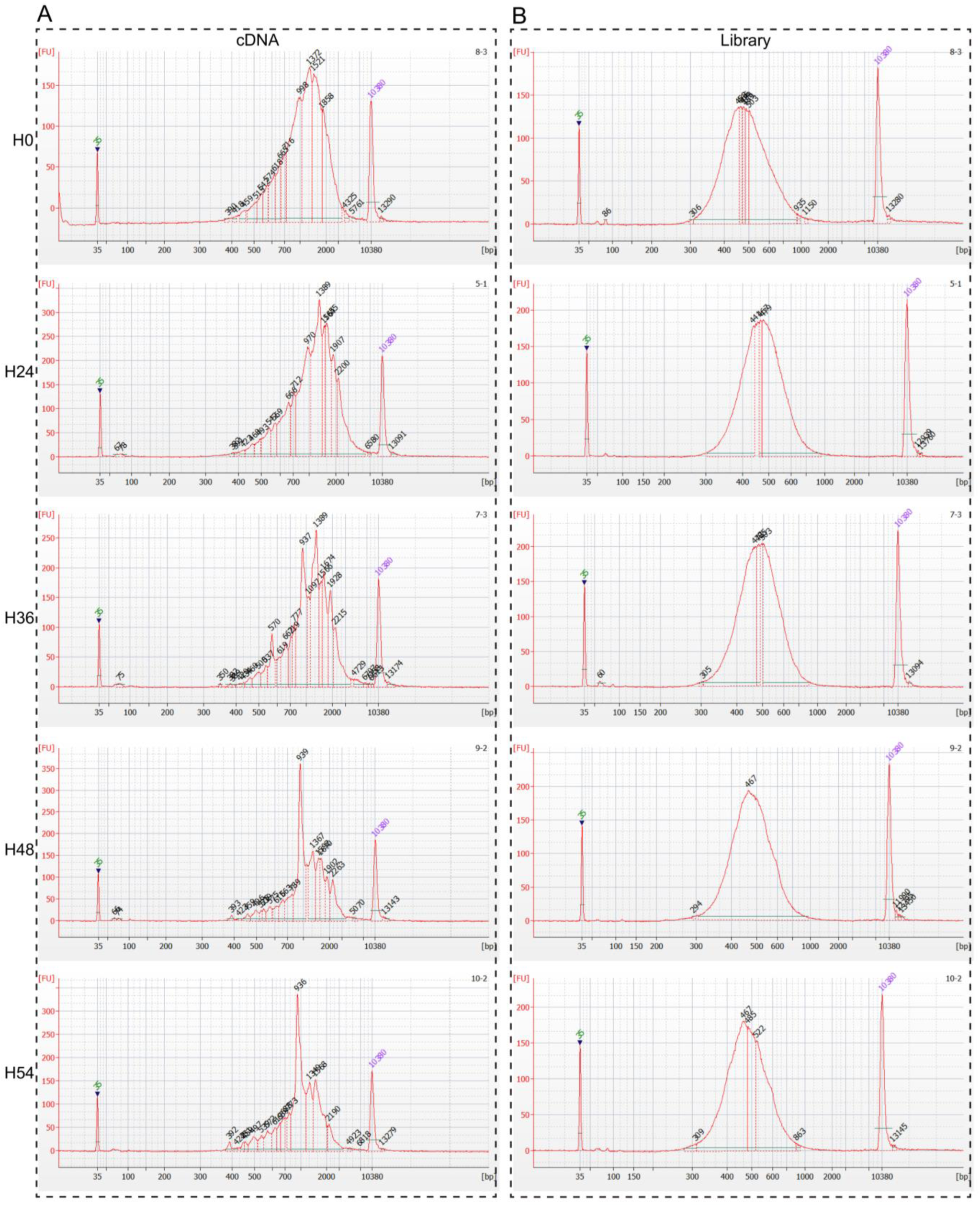
cDNA and library quality profiles. (**A, B**) The electrophoretogram of cDNA and library of five group samples respectively. FU, fluorescence unit.

**Figure S3.**
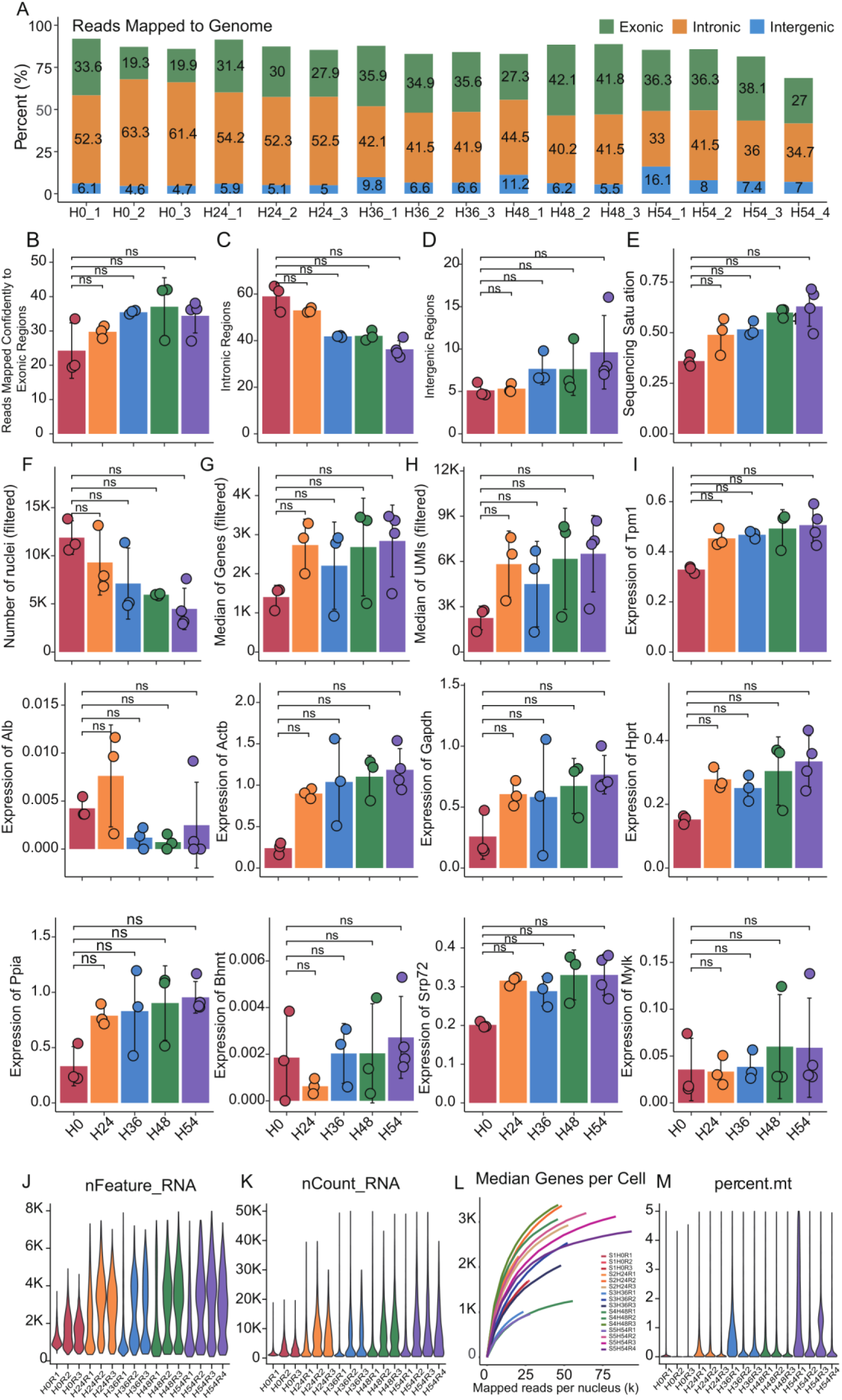
Quality assessment of sequencing data form five group nuclei. (A) Unique sequencing reads in genomic regions. The percentage of reads mapped to the genome, including exons, introns, and intergenic regions. (B-H) Gene exonic regions (B), intronic regions (C), intergenic regions (D), sequencing saturation (E), total number of nuclei detected (F), median genes (G) and median UMIs (H) at five PMI point. (I) Transcript levels of housekeeping genes among all PMI. (J, K) The violin plot shows the distribution of the number of genes (J) and transcripts (K) detected in each sample. (L) Gene sensitivity analysis of all samples; (M) The percentage of mitochondrial genes in all samples. n = 3 biologically independent mouse brain samples per group. Bars show mean ± s.d.

**Figure S4.**
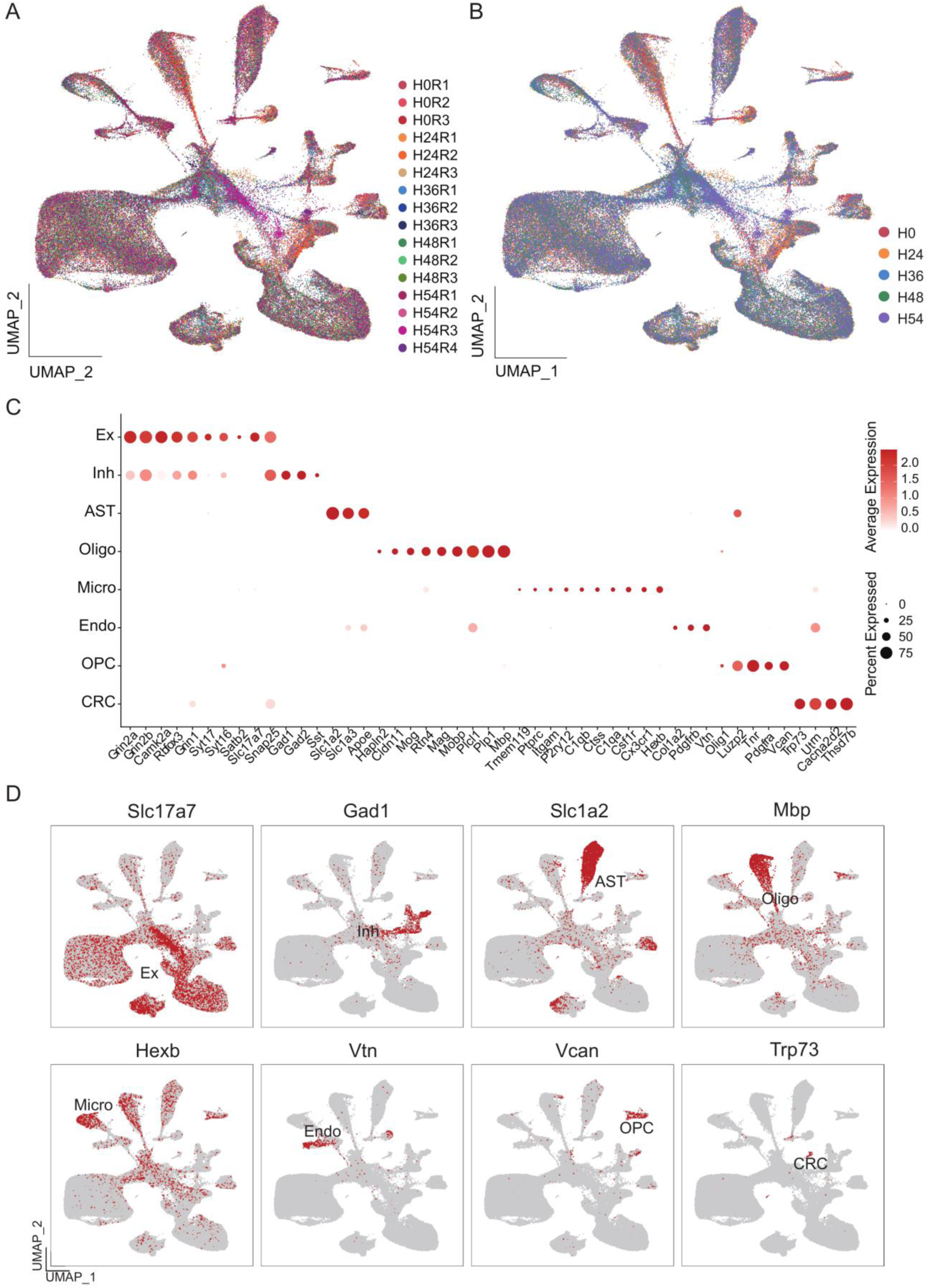
A cellular map of the mouse hippocampus of five PMI. (**A, B**) UMAP embedding of single nuclei RNA profiles from hippocampus of all samples (as in Fig. 1C), colored by samples (A) and groups (B). (**C**) Clusters and marker genes. Dot plot showing the expression level (color scale) and the percent of cells expressing (dot size) marker genes across all cells (rows). Cell types as in Fig. 1B. (**D**) Expression of marker genes across clusters. Colored by expression levels of marker genes: *Slc17a7* (excitatory neurons, Ex), *Gad1* (Inhibitory neurons, Ihn), *Slc1a2* (astrocytes, Ast), *Mbp* (oligodendrocytes, Oligo), *Hexb* (microglia, Micro), *Vtn* (endothelial, Endo), *Vcan* (Oligodendrocytes precursor cells, OPCs), *Trp73* (cajal-retzius cell,CRC).

**Figure S5.**
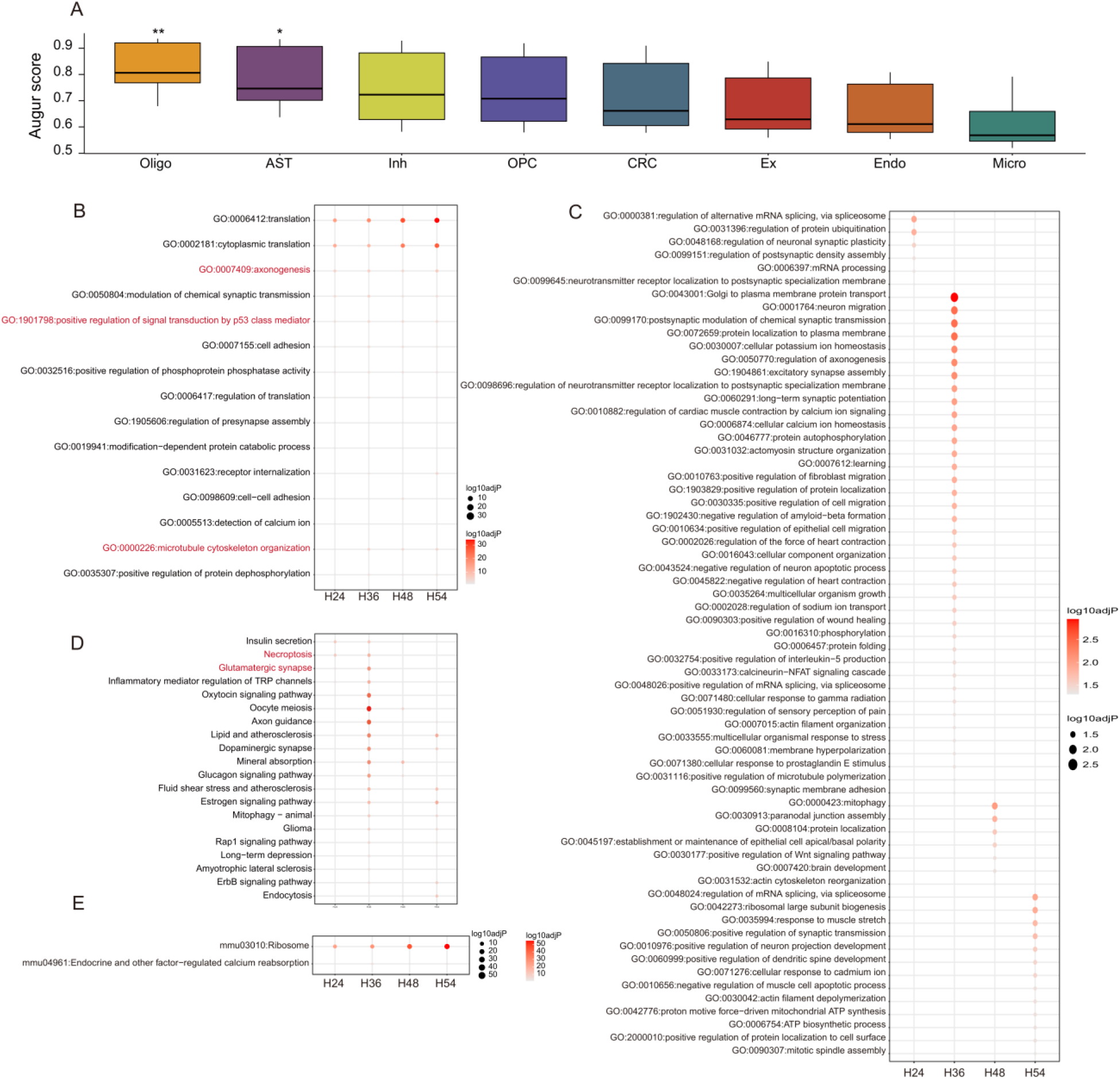
Cell-type-specific transcriptomic changes assessed by snRNA-seq across various ischaemia intervals. (**A**) Comparison of averaged Augur area under the curve (AUC) scores across the t-types indicating which cell type underwent the most transcriptomic changes. (**B, C**) *P* values of gene set enrichment of gene sets of specific cellular functions in Oligo cells. Shared functions (B) and PMI specific functions (C). (**D, E**) Shared (D) and PMI specific pathways (E) of Oligo gene set.

**Figure S6.**
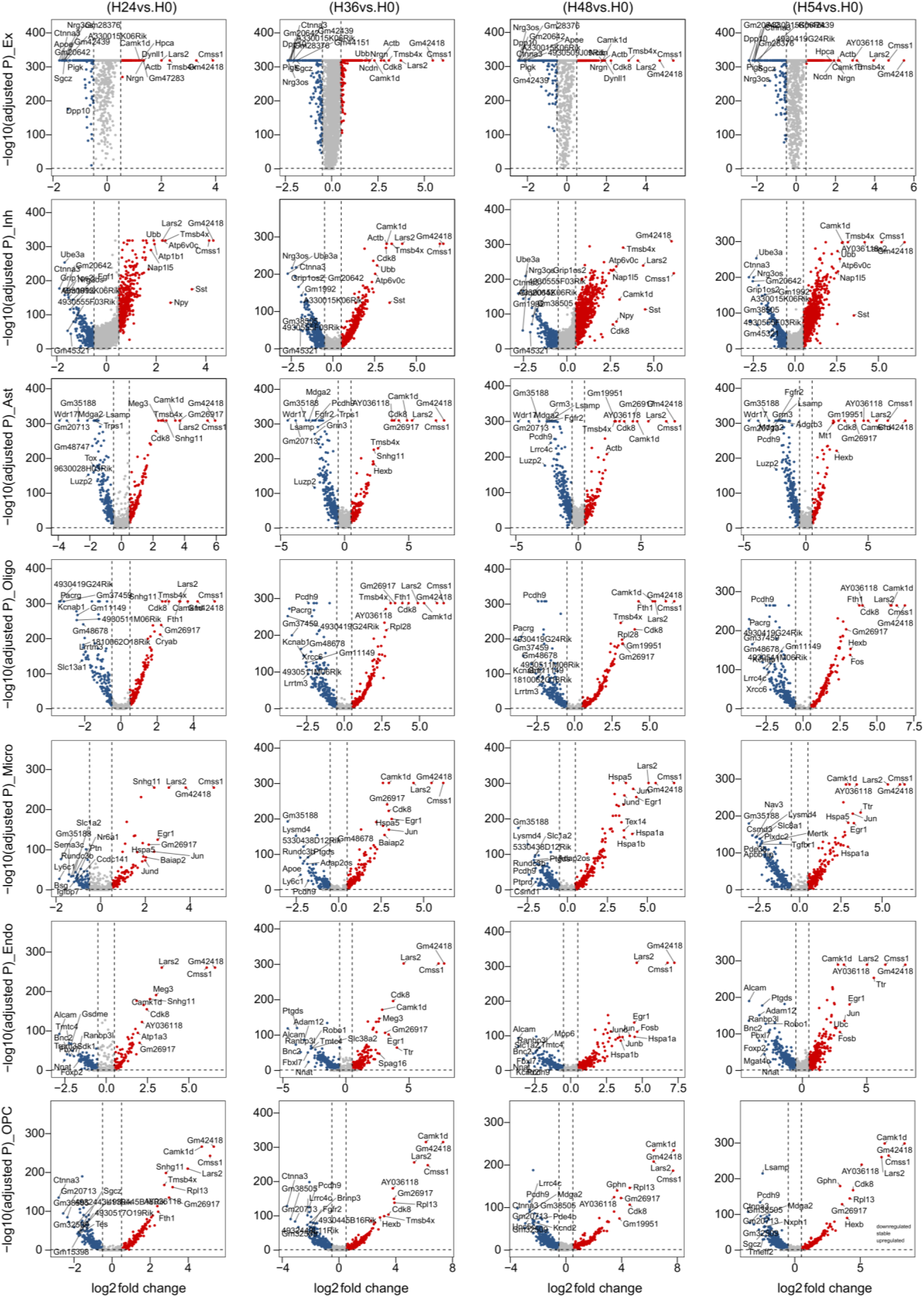
Transcriptional programs of all cell types and five PMI groups. Differential expression across all cell types. Volcano plots showing differentially expressed genes in each pair of states (n=3 animals. y-axis: -log adjusted hypergeometric p-value, following FDR multiple hypothesis correction, x-axis: average log fold change). Red and blue respectively indicated that the gene in the H24-H54 sample was significantly (*p* < 0.05)up and down regulated, and the grey was not significant (*p* > 0.05).

**Figure S7.**
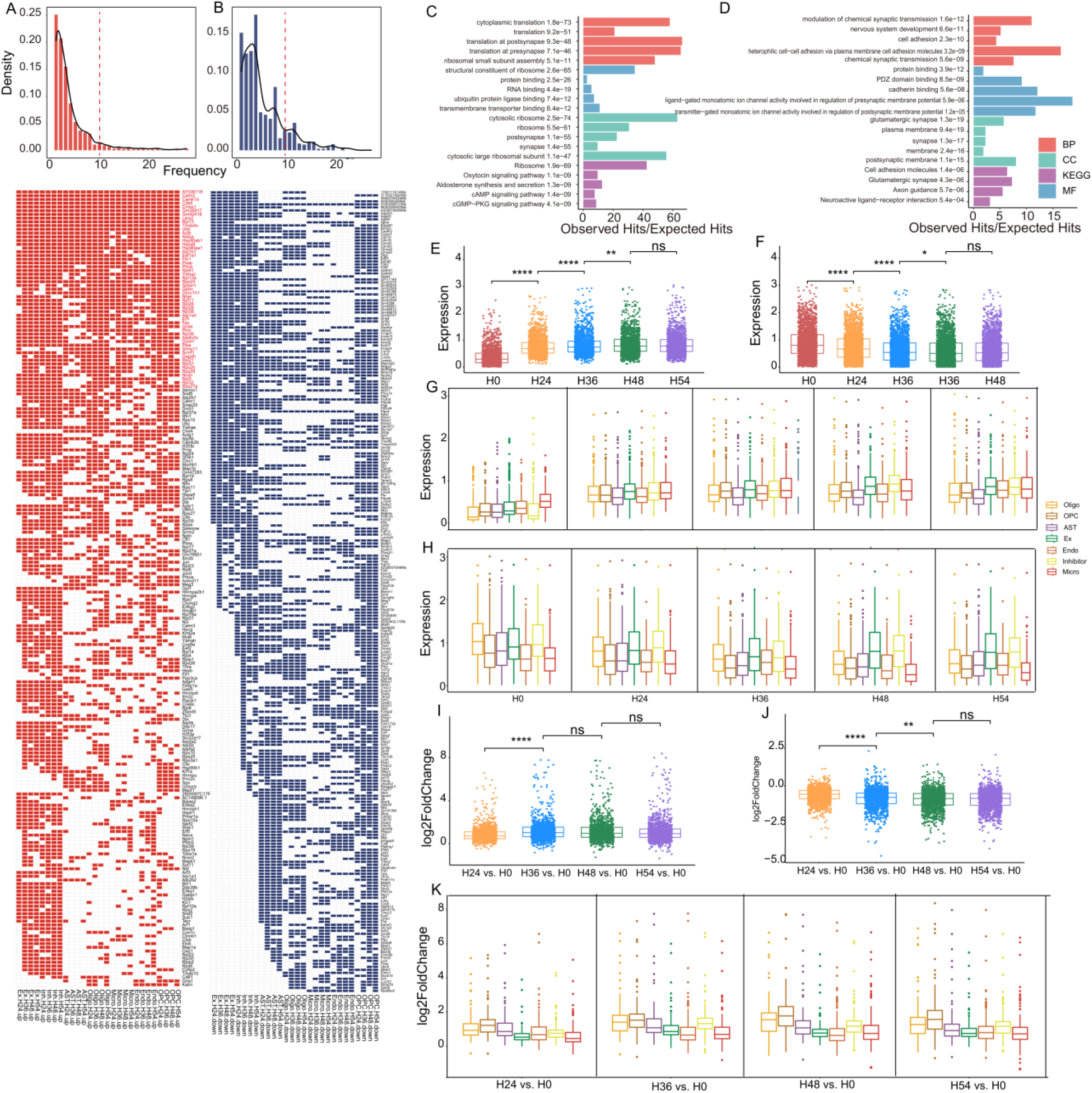
Identifies the up- and down-regulated genes by the difference expression gene analysis. (**A, B**) The distribution density chart of up- and down-regulated DEGs (top), and the shared DEGs in all cell types of five groups (bottom). (**C, D**) GO and KEGG analysis of A and B. (E, F)

**Figure S8.**
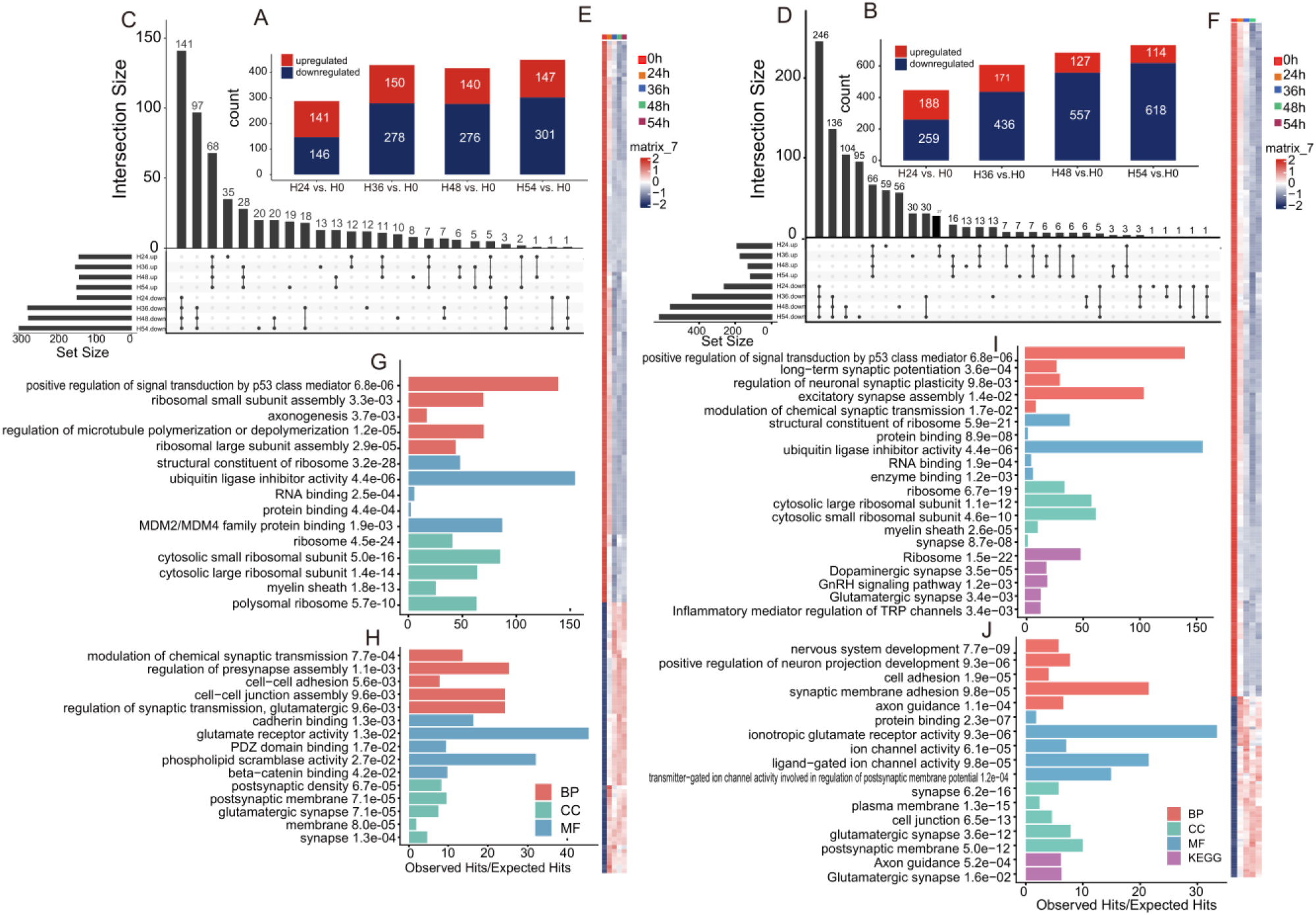
Differentially expressed genes analysis of in oligodendrocytes and OPCs. (**A, B**) H24-54 data for Oligo cells and OPCs and number of up-and down-regulated DEGs for H0. (**C, D**) Number of shared DEGs among all comparison groups. (**E, F**) Heatmap of co-up-regulated and co-down-regulated DEGs at five time points. Bar graph of (G, H) GO and KEGG analysis of co-up-regulated and down-regulated DEGs in Oligo cells. (**I, J**) Bar graph of GO and KEGG analysis of co-up-down-regulated DEGs in OPCs.

**Figure S9.**
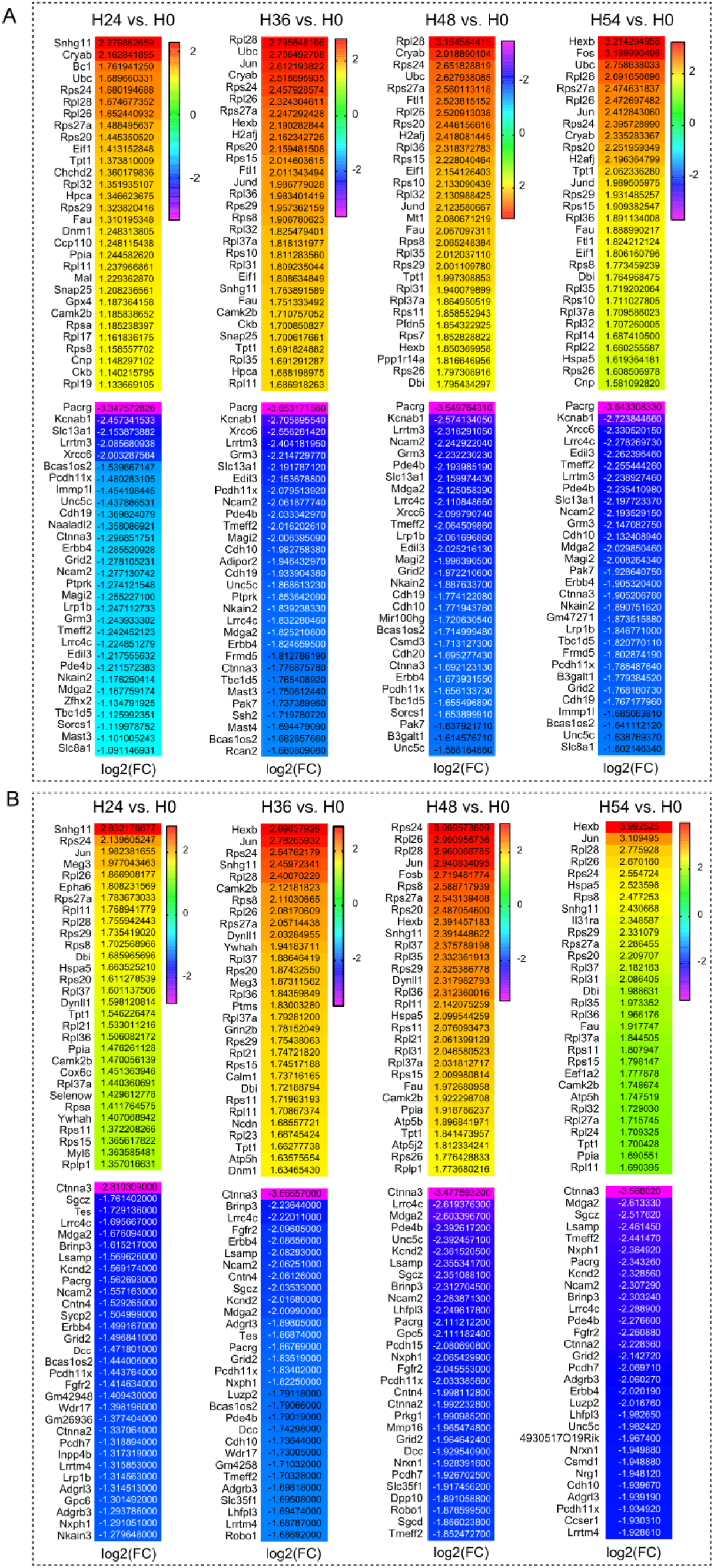
PMI causes up-and downregulation of genes. (**A, B**) Expression of the top genes with highest variance in Oligo (A) and OPCs (B).

**Figure S10.** Subcluster Analysis of excitatory neuronal cells. (**A**) Expression changes of specific highly expressed genes in Ex cells at five time points after death. (**B**) Volcano map of DEGs in Ex cells. (**C, D**) number of significantly up-regulated and down-regulated genes (C) and number of shared genes (D) in Ex cells. (**E**) GO functional analysis of down-regulated genes. (**F**) Clustering map of Ex subclusters (left), mapping in the original UMAP map (middle), and bubble map of specific expressed genes (right). (**G**) UMAP of Ex sub-cluster at five time points after death. (**H**) the proportion of Ex subgroup cells and their normalization results. (**I**) GO and KEGG analysis of Ex subcluster1. (**J**) Expression of genes involved in endogenous apoptosis, ferroptosis and necroptosis pathways in Ex subclusters.

**Figure S11.** GO and KEGG analysis of subclusters of excitatory neurons.

**Figure S12.**
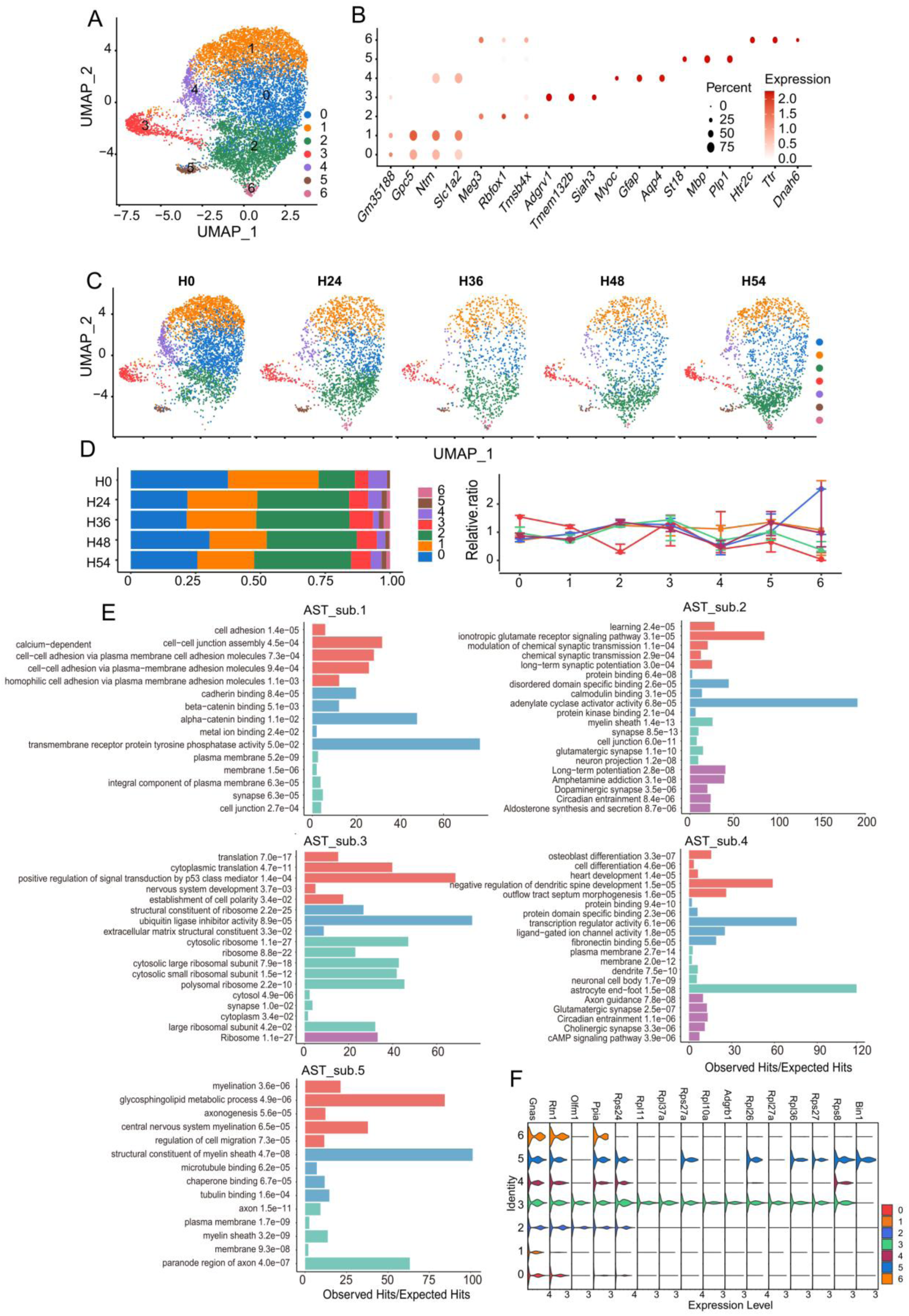
Subpopulations analysis of astrocytes. (**A**) Clustering map of AST subclusters, mapping in the original UMAP. (**B**) Bubble map of specific expressed genes. (**C**) UMAP of AST sub-cluster at five time points after death. (**D**) The proportion of AST subgroup cells and their normalization results. (**E**) GO and KEGG analysis of AST subclusters. (**F**) Expression of RB genes in AST subclusters.

**Figure S13.**
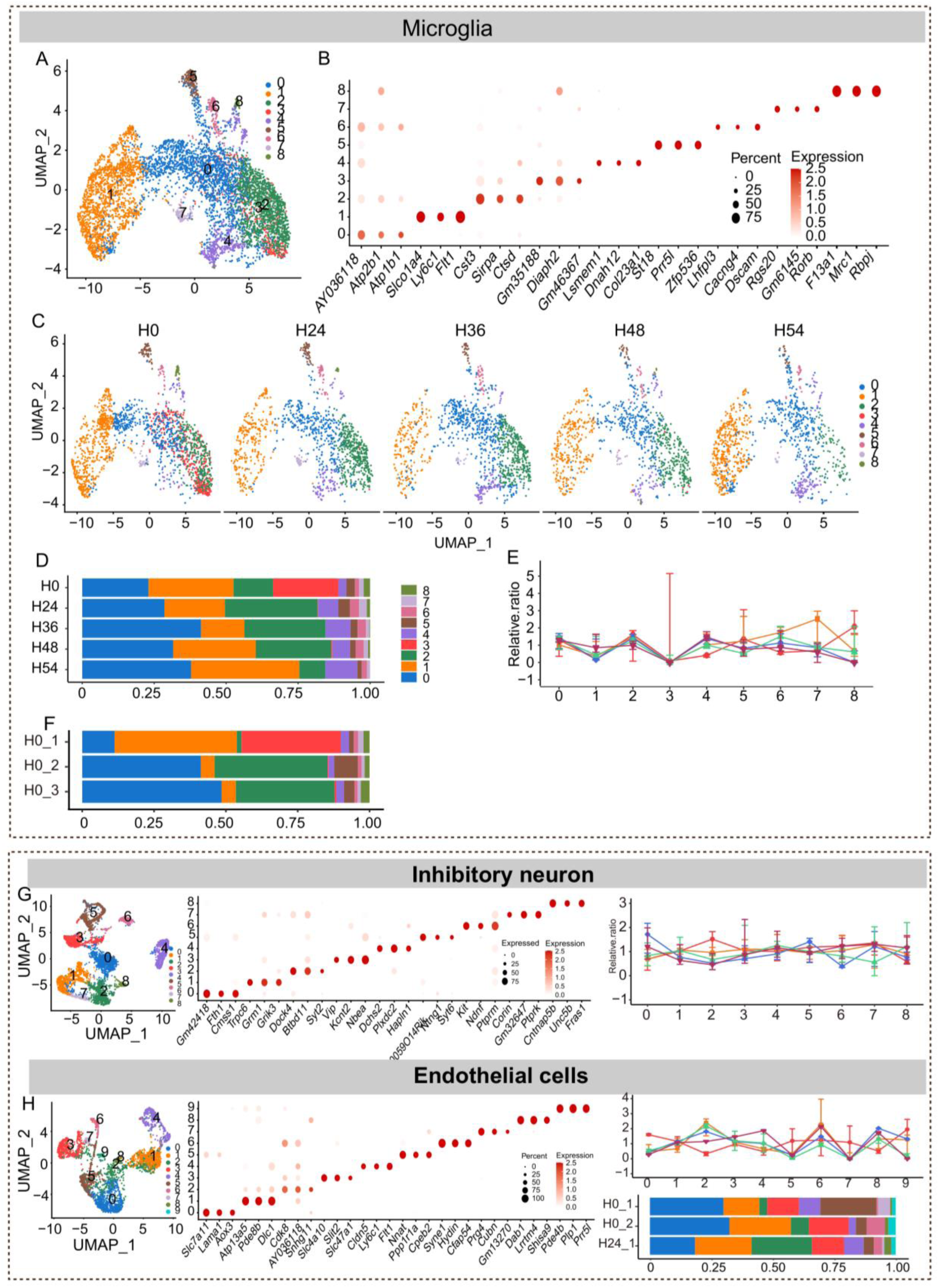
Subcluster analysis of microglia, inhibitory neurons, and endothelial cells.

**Figure S14.**
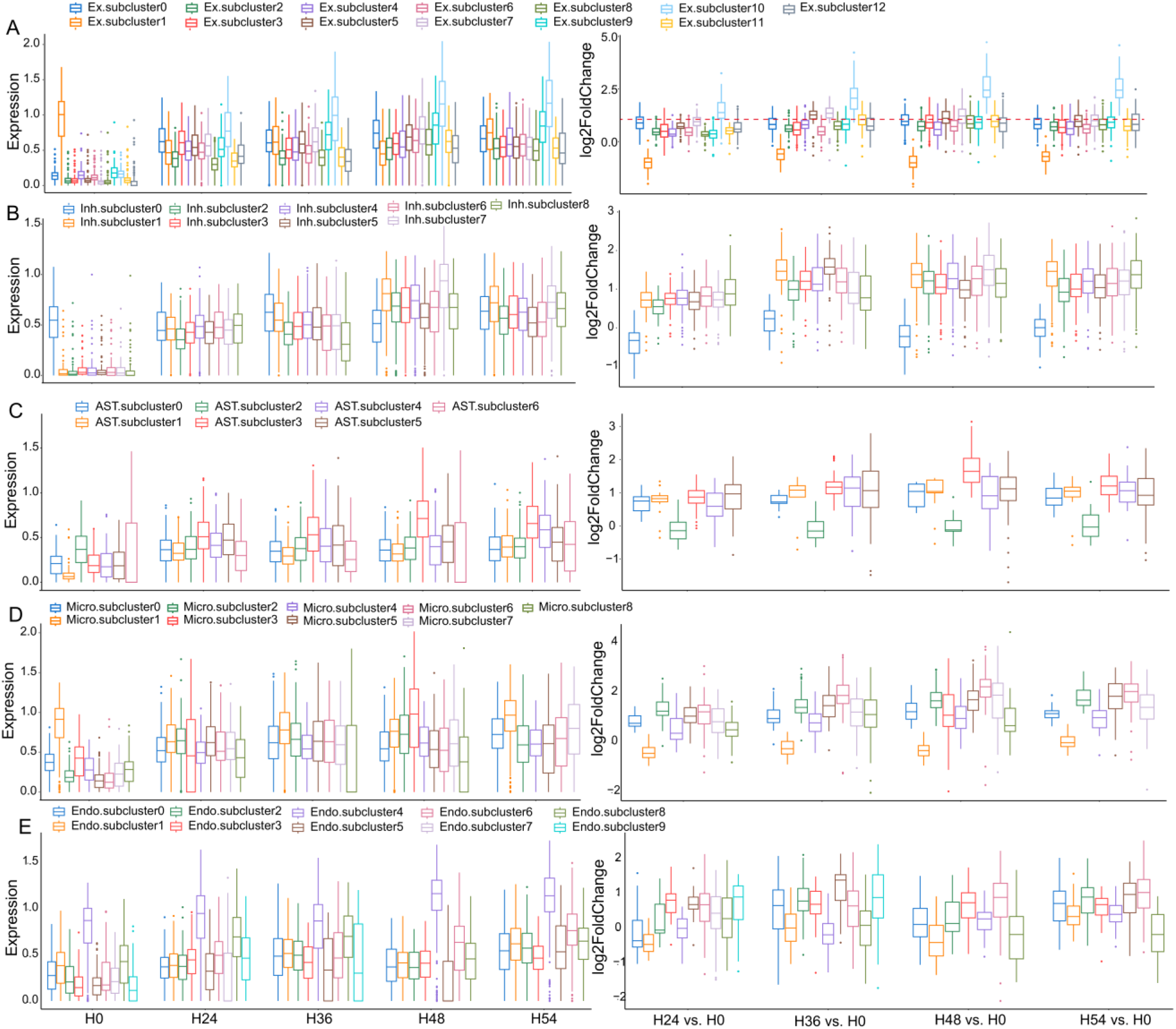
Expression Analysis of ribosomal genes in other subsets. From top to bottom, the expression changes (left) and fold change (right) of ribosomal genes in excitatory (A), inhibitory (B), astrocyte (C), microglia (D) and endothelial (E) subcluster cells are analyzed.

**Figure S15.**
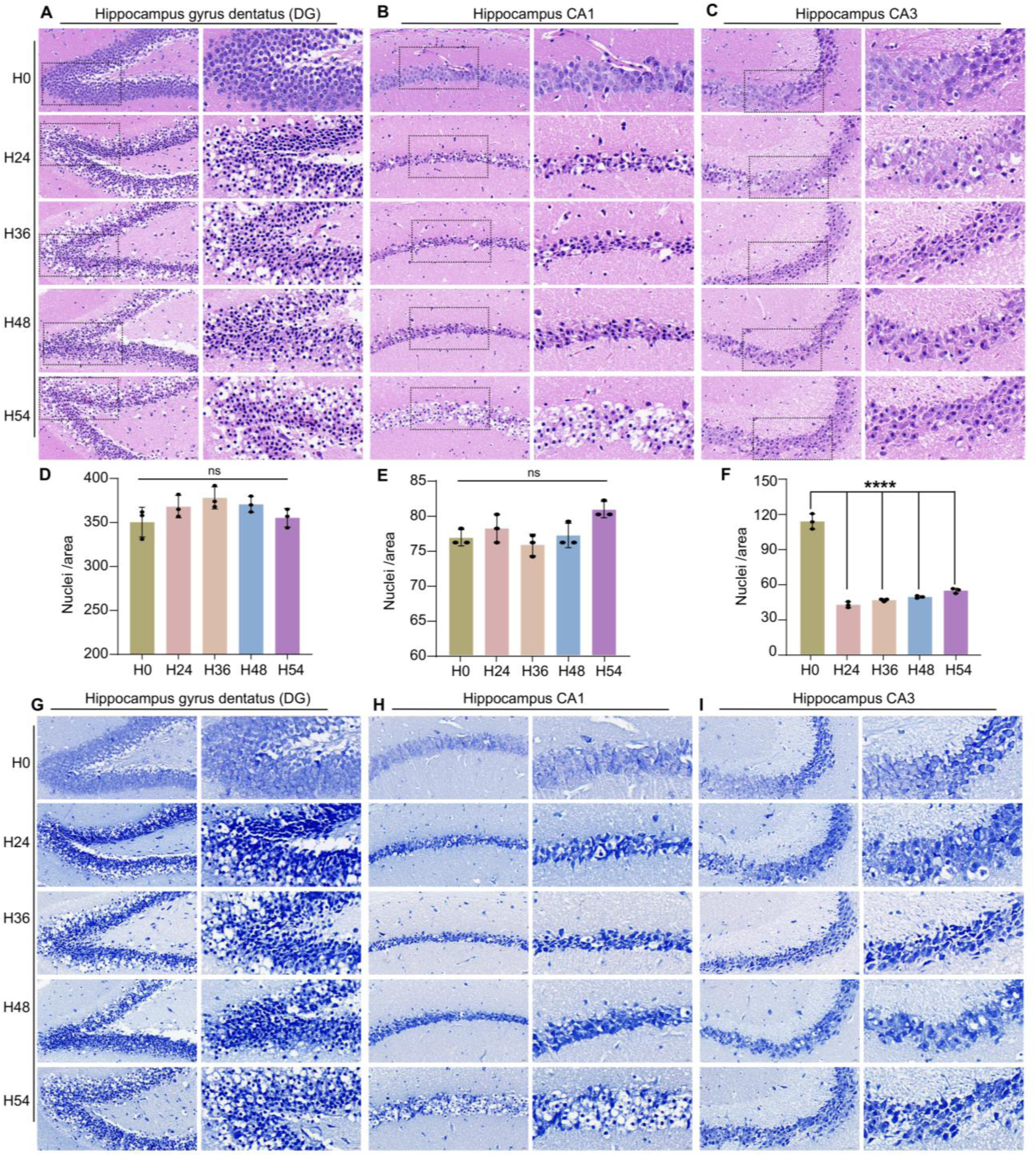
Analysis of tissue integrity across experimental conditions. (**A-C**) Representative images of H&E staining in the DG, CA1 and CA3 regions of hippocampus. (**D-F**) Quantification of the number of nuclei per standardized area in A-C between the experimental groups. (**G-H**) Representative images of Nissl staining of the DG, CA1 and CA3. n = 3. Scale bars, 20 μm (left) and 10 μm (right). Data presented are mean ± s.e.m. One-way ANOVA with post-hoc Dunnett’s adjustments was performed. For more detailed information on statistics and reproducibility, see methods. *P < 0.05, **P < 0.01, ***P < 0.001.

**Figure S16.**
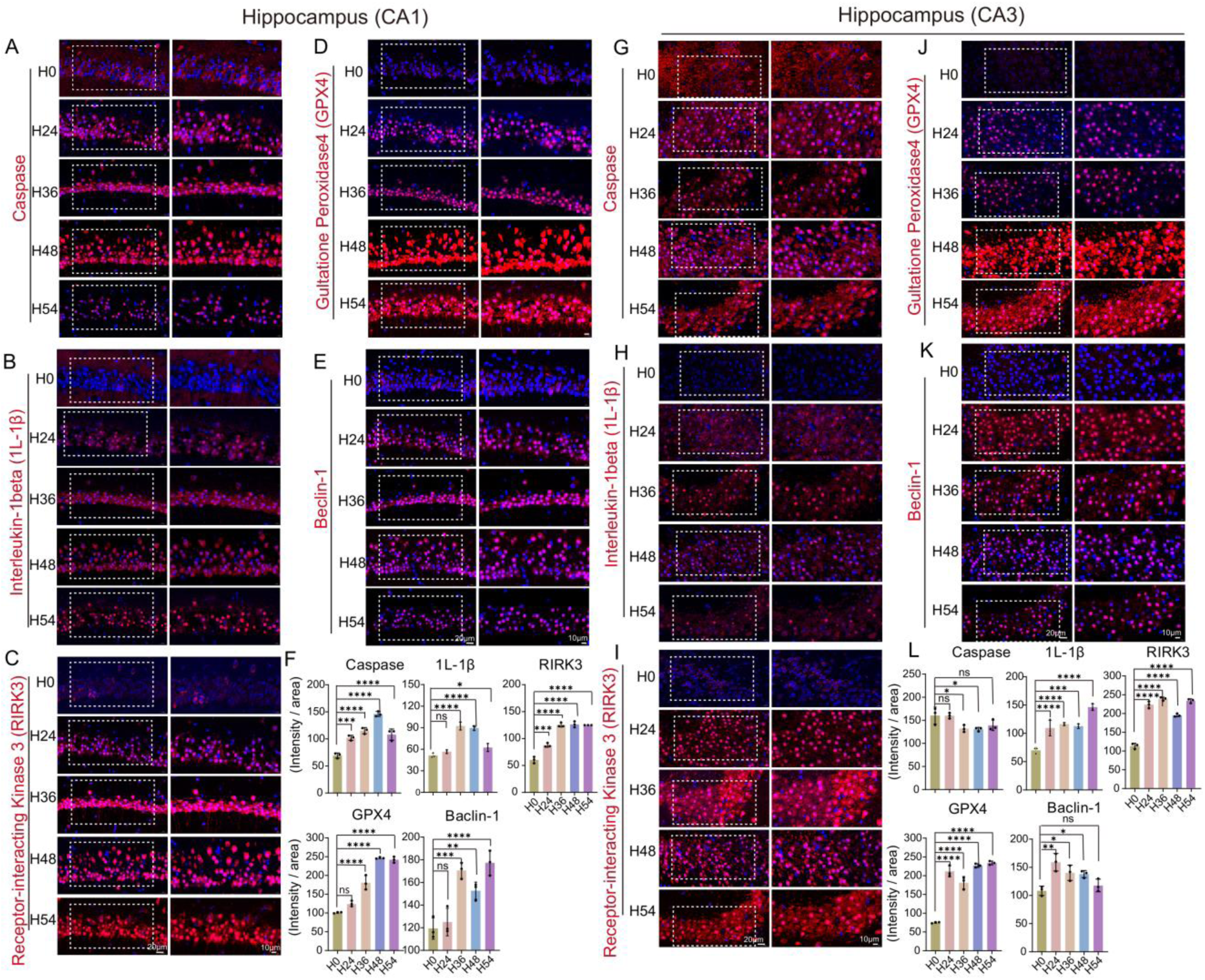
Analysis of cell death across experimental conditions. (**A-E**) Representative confocal images of immunofluorescent staining for activated caspase 3 (actCASP3), pyroptosis (IL-1β), necroptosis (RIPK3), ferroptosis (GPX4) and autophagy (Beclin-1) in CA1 of hippocampus, and co-stained with DAPI nuclear stain. (**F**) Quantification of actCASP3, IL-1β, RIPK3, GPX4 and Beclin-1 immunolabeling signal intensity in CA1. (**G-K**) Representative confocal images of immunofluorescent staining for activated caspase 3 (actCASP3), pyroptosis (IL- 1β), necroptosis (RIPK3), ferroptosis (GPX4) and autophagy (Beclin-1) in CA3 of hippocampus, and co-stained with DAPI nuclear stain. (**F**) Quantification of actCASP3, IL-1β, RIPK3, GPX4 and Beclin-1 immunolabeling signal intensity in CA3. n = 3. Scale bars, 20 μm (left) and 10 μm (right). Data presented are mean ± s.e.m. One-way ANOVA with post-hoc Dunnett’s adjustments was performed. For more detailed information on statistics and reproducibility, see methods. *P < 0.05, **P < 0.01, ***P < 0.001.

**Figure S17.**
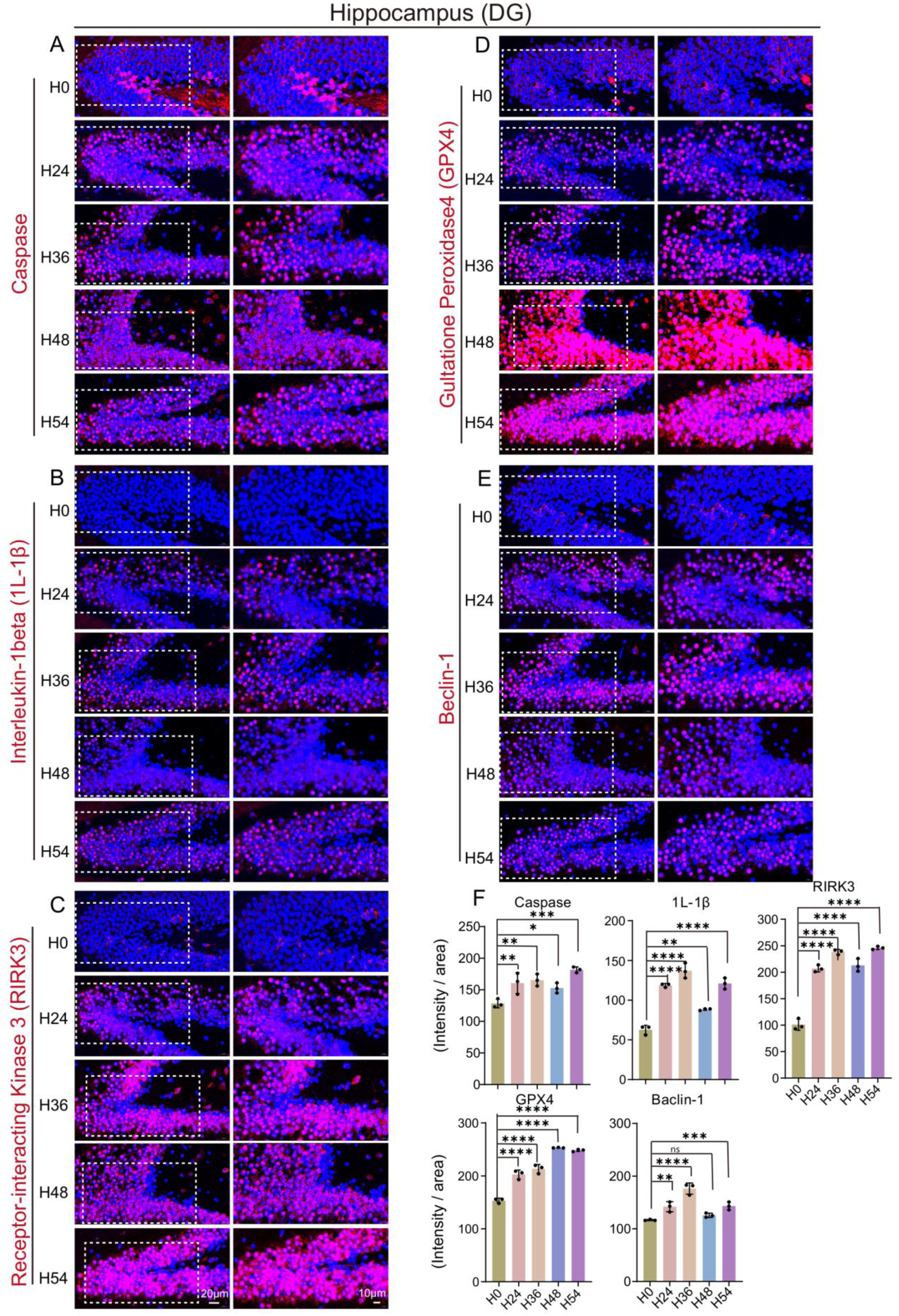
Analysis of cell death in DG across experimental conditions. (**A-E**) Representative confocal images of immunofluorescent staining for activated caspase 3 (actCASP3), pyroptosis (IL-1β), necroptosis (RIPK3), ferroptosis (GPX4) and autophagy (Beclin-1) in DG of hippocampus, and co-stained with DAPI nuclear stain. (**F**) Quantification of actCASP3, IL-1β, RIPK3, GPX4 and Beclin-1 immunolabeling signal intensity in DG. n = 3. Scale bars, 20 μm (left) and 10 μm (right). Data presented are mean ± s.e.m. One-way ANOVA with post-hoc Dunnett’ s adjustments was performed. For more detailed information on statistics and reproducibility, see methods. *P < 0.05, **P < 0.01, ***P < 0.001.

**Figure S18.**
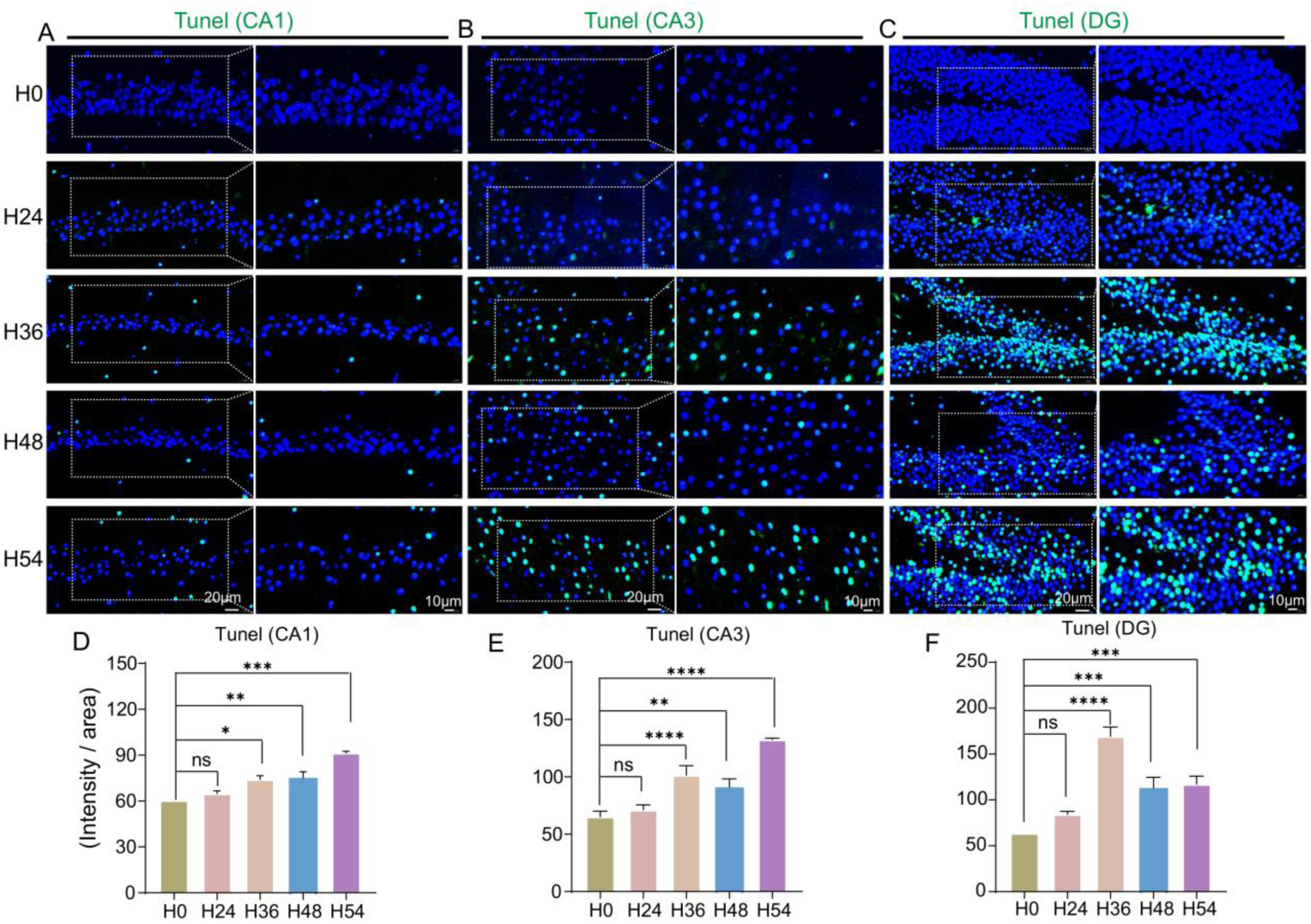
(**A-C**) The immunofluorescent images staining of TUNEL in CA1, CA3 and DG. (**D-F**) Normalized intensity of TUNEL signal in A-C. n = 3. Scale bars, 20 μm (left) and 10 μm (right). Data presented are mean ± s.e.m. One-way ANOVA with post-hoc Dunnett’s adjustments was performed. For more detailed information on statistics and reproducibility, see methods. *P < 0.05, **P < 0.01, ***P < 0.001.

## Reference

[1] Lee P, Chandel N S, Simon M C. Cellular adaptation to hypoxia through hypoxia inducible factors and beyond. Nat Rev Mol Cell Biol, 2020, 21: 268–283.

[2] Gonzalez-Herrera L, Valenzuela A, Marchal J A, et al. Studies on RNA integrity and gene expression in human myocardial tissue, pericardial fluid and blood, and its postmortem stability. Forensic Sci Int, 2013, 232: 218–28.

[3] Fordyce S L, Kampmann M-L, Doorn N L v, et al. Long-term RNA persistence in postmortem contexts. Investigative Genetics 2013, 4: 7–14.

[4] Partemi S, Berne P M, Batlle M, et al. Analysis of mRNA from human heart tissue and putative applications in forensic molecular pathology. Forensic Sci Int, 2010, 203: 99–105.

[5] Lee J, Hever A, Willhite D, et al. Effects of RNA degradation on gene expression analysis of human postmortem tissues. FASEB J, 2005, 19: 1356–1358.

[6] Fitzpatrick R, Casey O M, Morris D, et al. Postmortem stability of RNA isolated from bovine reproductive tissues. Biochimica et Biophysica Acta (BBA) - Gene Structure and Expression, 2002,

[7] Daniele S G, Trummer G, Hossmann K A, et al. Brain vulnerability and viability after ischaemia. Nat Rev Neurosci, 2021, 22: 553–572.

8. Kirino T. Delayed Neuronal Death In The Gerbil Hippocampus. Brain Research, 1982, 239: 57–69.

[9] Kirino T, Sano K. Fine structural nature of delayed neuronal death following ischemia in the gerbil hippocampus. Acta Neuropathologica, 1984, 62: 209–218.

[10] Dewar D, Underhill S M, Goldberg M P. Oligodendrocytes and ischemic brain injury. J Cereb Blood Flow Metab, 2003, 23: 263–274.

[11] Benjamín S, Olof E, Joakim L, et al. Sequencing Degraded RNA Addressed by 3’ Tag Counting. PLoS ONE, 2014, 9: e91851.

[12] Feng H, Zhang X, Zhang C. mRIN for direct assessment of genome-wide and gene-specific mRNA integrity from large-scale RNA-sequencing data. Nature Communications, 2015, 6: 7816.

[13] Romero I G, Pai A A, Tung J, et al. RNA-seq: impact of RNA degradation on transcript quantification. Bmc Biology, 2014, 12: 42–45.

[14] Irene Gallego Romero, Athma A Pai, Jenny Tung, et al. RNA-seq: impact of RNA degradation on transcript quantification. BMC Biology 2014, 12: 42–55.

15. Guo Y, Ma J, Dang K, et al. Transcriptomic profiling of nuclei from PFA-fixed and FFPE brain tissues. bioRxiv preprint doi: 10.1101/2023.04.13.536693, 2023,

[16] Morgan J T, Fink G R, Bartel D P. Excised linear introns regulate growth in yeast. Nature, 2018, 565: 606.

[17] Parenteau J, Maignon L, Berthoumieux M, et al. Introns are mediators of cell response to starvation. Nature, 2019, 565: 612–617.

[18] Dachet F, Brown J B, Valyi-Nagy T, et al. Selective time-dependent changes in activity and cell-specific gene expression in human postmortem brain. Sci Rep, 2021, 11: 6078.

[19] Guo Y, Ma J, Li Z, et al. Transcriptomic profiling of nuclei from paraformaldehyde-fixed and formalin-fixed paraffin-embedded brain tissues. Analytica Chimica Acta, 2023, 1281: 341861.

[20] Guo Y, Ma J, Huang H, et al. Defining Specific Cell States of MPTP-Induced Parkinson’s Disease by Single-Nucleus RNA Sequencing. Int. J. Mol. Sci., 2022, 23: 10774–10794.

[21] Zhou Y, Song W M, Andhey P S, et al. Human and mouse single-nucleus transcriptomics reveal TREM2- dependent and TREM2-independent cellular responses in Alzheimer’s disease. Nat. Med., 2020, 26: 131–142.

[22] Habib N, McCabe C, Medina S, et al. Disease-associated astrocytes in Alzheimer’s disease and aging. Nat. Neurosci., 2020, 23: 701–706.

23. Skinnider M A, Squair J W, Kathe C, et al. Cell type prioritization in single-cell data. Nature Publishing Group, 2021, 39: 30–34.

[24] Andrijevic D, Vrselja Z, Lysyy T, et al. Cellular recovery after prolonged warm ischaemia of the whole body. Nature, 2022, 608: 405–412.

[25] Brumwell A, Fell L, Obress L, et al. Hypoxia influences polysome distribution of human ribosomal protein S12 and alternative splicing of ribosomal protein mRNAs. RNA (New York, N.Y.), 2020, 26: 361–371.

[26] Xie J Q, Lu Y P, Sun H L, et al. Sex Difference of Ribosome in Stroke-Induced Peripheral Immunosuppression by Integrated Bioinformatics Analysis. Biomed Res Int, 2020, 2020: 3650935.

27. Chen Y, Fan Z, Wu Q. Dexmedetomidine improves oxygen-glucose deprivation/reoxygenation (OGD/R) - induced neurological injury through regulating SNHG11/miR-324-3p/VEGFA axis.

[28] Wang Z, Liu L, Guo X, et al. microRNA-1236-3p Regulates DDP Resistance in Lung Cancer Cells. Open Med (Wars), 2019,

[29] Taylor J M, Brody K M, Lockhart P J. Parkin Co-Regulated Gene is involved in aggresome formation and autophagy in response to proteasomal impairment. Experimental Cell Research, 2012, 318: 2059–2070.

[30] Bladen C L, Navarre S, Dynan W S, et al. Expression of the Ku70 subunit (XRCC6) and protection from low dose ionizing radiation during zebrafish embryogenesis. Neuroscience Letters, 2007, 422: 97–102.

[31] Park, Chaehwa. Maspin increases Ku70 acetylation and Bax-mediated cell death in cancer cells. International Journal of Molecular Medicine, 2012,

[32] Lee S H, Moon S J, Shin J B, et al. The Effect of the Water Extract of Angelica Sinens on Gliosis Repression of Astrocyte after Hypoxic injury. 29, 2008, 1:

[33] Frauke, Seehusen, Kirsten, et al. Axonopathy in the Central Nervous System Is the Hallmark of Mice with a Novel Intragenic Null Mutation of Dystonin. Genetics, 2016, 204: 191.

[34] Shen G, Hu S, Zhao Z, et al. C-Type Natriuretic Peptide Ameliorates Vascular Injury and Improves Neurological Outcomes in Neonatal Hypoxic-Ischemic Brain Injury in Mice. Multidisciplinary Digital Publishing Institute, 2021, 22: 8966.

[35] Riedhammer K M, Stockler S, Ploski R, et al. De novo stop-loss variants in CLDN11 cause hypomyelinating leukodystrophy (vol 144, pg 411, 2020). Brain: A journal of neurology, 2021, 144: e48.

[36] Li Z, Qian R, Zhang J, et al. Lipoma HMGIC fusion partner-like 3 (LHFPL3) promotes proliferation, migration and epithelial-mesenchymal transitions in human glioma cells. International Journal of Clinical and Experimental Pathology, 2017, 10: 5471–5479.

[37] Shin, J.-G., Kim, et al. Putative association of GPC5 polymorphism with the risk of inflammatory demyelinating diseases. Journal of the Neurological Sciences Official Bulletin of the World Federation of Neurology, 2013, 335: 82–88.

[38] He S, Li Y, Shi X, et al. DNA methylation landscape reveals LIN7A as a decitabine-responsive marker in patients with t(8;21) acute myeloid leukemia. Clinical Epigenetics, 2023, 15:

[39] Jeong-Min K, Kyu-Hwa L, Yeo-Jin J, et al. Identification of Genes Related to Parkinson’s Disease Using Expressed Sequence Tags. DNA Research,13,6(2007-1-8), 2007, 275.

[40] Wang, L. Cell-Based Screening and Validation of Human Novel Genes Associated with Cell Viability. Journal of Biomolecular Screening, 2006, 11: 369.

[41] Hsia C C W, Schmitz A, Lambertz M, et al. Evolution of Air Breathing: Oxygen Homeostasis and the Transitions from Water to Land and Sky. John Wiley & Sons, Inc., 2013,

[42] Matsuda T. Importance of experimental information (metadata) for archived sequence data: case of specific gene bias due to lag time between sample harvest and RNA protection in RNA sequencing. PeerJ, 2021, 9: e11875.

[43] Lowry O H, Passonneau J V, Hasselberger F X, et al. Effect of Ischemia on Known Substrates and Cofactors of the Glycolytic Pathway in Brain. Journal of Biological Chemistry, 1964, 239: 18–30.

[44] D M A F A, D N A E C, D I I T, et al. Effect of seasonal variation in ambient temperature on RNA quality of breast cancer tissue in a remote biobank setting - ScienceDirect. Experimental and Molecular Pathology, 112:

[45] Auer H, Mobley J, Ayers L, et al. The effects of frozen tissue storage conditions on the integrity of RNA and protein. Biotechnic & Histochemistry, 2014, 89: 518–528.

[46] Shen Y, Li R, Tian F, et al. Impact of RNA integrity and blood sample storage conditions on the gene expression analysis. Onco Targets Ther, 2018, 11: 3573–3581.

[47] Kono N, Nakamura H, Ito Y, et al. Evaluation of the impact of RNA preservation methods of spiders for de novo transcriptome assembly. Molecular Ecology Resources, 2015, 16: 662–672.

[48] Hu H, Li X. Transcriptional regulation in eukaryotic ribosomal protein genes. Genomics, 2007, 90: 421–423.

[49] Fisher E M C, Beer-Romero P, Brown L G, et al. Homologous ribosomal protein genes on the human X and Y chromosomes: escape from X inactivation and possible implications for Turner syndrome. Cell, 1990, 63: 1205–1218.

[50] Radek, Cmejla, Jana, et al. Ribosomal protein S17 gene (RPS17) is mutated in Diamond-Blackfan anemia. Human Mutation, 2007, 28: 1178–1182.

[51] Uechi T, Tanaka T, Kenmochi N. A complete map of the human ribosomal protein genes: assignment of 80 genes to the cytogenetic map and implications for human disorders. Genomics, 2001, 72: 223–230.

[52] Wang H, Wang B, Normoyle K P, et al. Brain temperature and its fundamental properties: a review for clinical neuroscientists. Front Neurosci, 2014, 8: 307.

[53] Ruonan J, Feng L, Qianghua X. Differential expression analysis of the ribosomal protein gene family in zebrafish gills under hypoxia stress. JOURNAL OF SHANGHAl OCEAN UNIVERSITY, 2022, 31: 318–327.

[54] Audinat S C. Cardiac arrest in rodents: maximal duration compatible with a recovery of neuronal activity. Proceedings of the National Academy of Sciences of the United States of America, 1998, 95:

[55] Onorati M, Li Z, Liu F, et al. Zika Virus Disrupts Phospho-TBK1 Localization and Mitosis in Human Neuroepithelial Stem Cells and Radial Glia - ScienceDirect. Cell Reports, 2016, 16: 2576–2592.

56. Molecular and cellular reorganization of neural circuits in the human lineage. Science, 2017, 358: 1027–1032.

[57] Xiong Z G, Zhu X M, Chu X P, et al. Neuroprotection in ischemia: blocking calcium-permeable acid-sensing ion channels. Cell, 2004, 118: 687–698.

[58] Goldberg M, Choi D. Combined oxygen and glucose deprivation in cortical cell culture: calcium-dependent and calcium-independent mechanisms of neuronal injury. Journal of Neuroscience, 1993, 13: 3510.

[59] Husnia, Marrif, Bernhard, et al. Astrocytes respond to hypoxia by increasing glycolytic capacity. Journal of Neuroscience Research, 1999,

[60] Garcia J H, Yoshida Y, Chen H, et al. Progression from ischemic injury to infarct following middle cerebral artery occlusion in the rat. American Journal of Pathology, 1993, 142: 623–35.

61. Baldea I, Teacoe I, Olteanu D E, et al. Effects of different hypoxia degrees on endothelial cell cultures - time course study. Mechanisms of Ageing & Development, 2017, S0047637417301215.

[62] Isidre F, Gabriel S, Thomas A, et al. Brain protein preservation largely depends on the postmortem storage temperature: implications for study of proteins in human neurologic diseases and management of brain banks: a BrainNet Europe Study. J Neuropathol Exp Neurol, 2007, 35–46.

[63] Smart J L, Kaliszan M. The post mortem temperature plateau and its role in the estimation of time of death. A review. Legal Medicine, 2012, 14: 55–62.

